# Thoughts and cortical responses are shaped by the type of change at event boundaries

**DOI:** 10.64898/2026.10.12.758535

**Authors:** Samuel Ketcheson, Raven S. Wallace, Silvia S. Zhou, Samyogita Hardikar, Bridget Mulholland, Louis Chitiz, Jason Gallivan, Robert Leech, Jonathan Smallwood, Elizabeth Jefferies, Jeffrey D. Wammes

## Abstract

Continuous experience is divided into discrete and meaningful events, but different kinds of event transitions may shape thought and brain activity in different ways. We examined how perceived event boundaries relate to visual, auditory, and semantic features, ongoing thought, and large-scale cortical states during movie-watching. We combined three independent samples who viewed the same eight movie clips, aligning these measures to a shared stimulus timeline: One marked event boundaries, one reported ongoing thought, and one was scanned with fMRI. Whole-brain spatial correlations showed that event boundaries correlated positively with visual change and negatively with auditory change, but not with any thought pattern. Time-resolved analyses showed that boundaries were associated with a reduction in distracted thoughts and brain states positioned toward visual processing. Clustering boundaries by movie feature change profile identified *visual, semantic, interruption, and narrative* boundary types. All four types showed a shift toward visual processing in the brain, but their other associations differed. Visual boundaries were associated with less distraction, semantic boundaries with thoughts related to past experience and prior knowledge, and narrative boundaries with more detailed and deliberate thoughts. Visual and semantic boundaries also showed distinct shifts in brain state that were not apparent in the other two types. These findings suggest that event boundaries mark a set of state shifts in ongoing experience, with a shared reorientation toward visual processing and changes in thought and brain state that vary with the type of transition.

## Introduction

As we navigate the world, subjective experience washes over us continuously, but people do not represent experience as an unstructured stream. Instead, they parse otherwise continuous activity into discrete, meaningful events (Baldassano et al., 2017), with considerable agreement across individuals (Zacks & Swallow, 2007). Event Segmentation Theory claims this is driven by event models, internal representations of the current event that are held in working memory to contend with constantly changing situations. When an event model’s predictions fail, sensory input is combined with prior knowledge to update the model, and a boundary is experienced (Zacks & Swallow, 2007). These established boundaries constrain our ongoing experience in a way that determines how they are stored in (Silva et al., 2019), structured, and later retrieved from memory (Flores et al., 2017; Gernsbacher, 1985; Kurby & Zacks, 2022), to the point where alignment with consensus predicts successful memory (Sargent et al., 2013). While often studied through where event boundaries fall, how the brain navigates the transition, and the consequent memory effects, event segmentation permeates our daily life (Zacks et al., 2010), meaning events likely go beyond what memories are formed, and also organize what occupies a person’s mind from moment to moment.

Event structure can be read out from brain activity. During movie-watching, patterns of cortical activity remain relatively stable within events, transitioning brain states at moments identified by observers as boundaries (Baldassano et al., 2017; Geerligs et al., 2022). The timescale of the events follows a cortical hierarchy, with short events represented in sensory regions and longer ones in higher-order association cortex (Baldassano et al., 2017; Geerligs et al., 2022). Changing brain states most robustly align with identified boundaries when they involve larger changes or span multiple brain regions (Geerligs et al., 2022). While the above establishes that there are consensus events, and that these coincide with changes in brain states and memory, events are variable (Cutting et al., 2012; Cutting, 2014; Kurby & Zacks, 2008; Zacks et al., 2010), and it is not obvious what gives rise to any particular event boundary.

Boundaries are not uniform occurrences; different kinds may arise from distinct changes in the unfolding environment, and the same sort of change might not always be detected as a boundary (Cutting et al., 2012; Kurby & Zacks, 2008). Even the prediction errors elicited by these changes can be driven by either episodic or schematic knowledge, with dissociable neural signatures (Varga et al., 2025). In movies, boundaries may coincide with abrupt perceptual changes (e.g. new location, soundtrack change), or higher-level conceptual changes (change in goals, narrative or dialogue). Reflecting this diversity, events have been profitably cross-sectioned by coding changes in character, location or time (Cutting, 2014; Zacks et al., 2010), but pretrained neural networks can directly quantify different sources of change continuously, yielding separate time courses for visual, auditory, and semantic change within the same movie. This makes it possible to identify which aspects of the changing environment are most strongly associated with boundary perception and the corresponding neural state transitions. More critically, if there are fundamentally different boundary types, then pooling their analyses will average over interesting variation in how events constrain cognitive and neural signatures.

A missing element in understanding these event transitions is measuring experience itself. The relationship between stimulus changes and event boundaries may depend on how viewers *think* during the unfolding narrative. This is an unexplored facet of event cognition, which is made especially important given the theory: If event models integrate incoming information with prior knowledge (Zacks & Swallow, 2007), then whether viewers’ thoughts are typically distracted, self-referential, affective, or related to memory during a particular moment will determine the structure of events. The above are just a few of the facets of ongoing thought (Smallwood et al., 2021) that fluctuate dynamically over time (Jääskeläinen et al., 2021; Konu et al., 2021; Saarimäki, 2021; Smallwood et al., 2021; Wallace et al., 2025; Zhang et al., 2021). Investigating this requires a method for characterizing ongoing experience, but traditionally, the dynamic richness of ongoing thought has been reduced to a single dimension, such as task focus, intentionality, or thought content (Weinstein, 2018; Kane et al., 2021). Multidimensional experience sampling (mDES) instead assesses sixteen dimensions of experience at each thought probe (e.g. self-focus, perceptual detail, social cognition; Smallwood et al., 2016; Smallwood et al., 2021), allowing the quantification of low-dimensional patterns of thought that predict behavioural performance, clinical traits, and brain organization (Chitiz et al., 2025; Mulholland et al., 2025; Smallwood et al., 2016; Smallwood et al., 2021; Wallace et al., 2025; Zhou et al., 2025). While movie-watching provides a naturalistic experience to measure ongoing cognition, frequent probe interruptions would break immersion, and disrupt the continuous nature of brain activity. Therefore, thought and brain data cannot be sampled from the same participants. Prior work has solved this problem by using uninterrupted fMRI data from one sample, then collecting infrequent thought probes from another sample outside the scanner to produce a group-level time course of thoughts that can be temporally mapped to the same stimulus, and in turn, the brain (Wallace et al., 2025). This approach therefore provides a structured framework for quantifying the content and dynamics of ongoing cognition, and how they relate to changing brain states around event boundaries.

Relating stimulus features, ongoing thought, event boundaries, and neural activity requires a common representational framework. Functional gradients solve this by describing cortical organization along a small number of continuous axes, the first extending from primary sensory and motor areas to association regions within the default mode network (Margulies et al., 2016). Gradients allow a complementary look at both the spatial organization and temporal dynamics of brain activity. First, by correlating an aggregate activity map with each gradient, we can characterize where there is sensitivity to a given feature or thought dimension in cortex, ignoring temporal fluctuation. Second, projecting moment-to-moment activity onto the same gradients asks whether large-scale activity patterns change over event boundaries, instead collapsing across space. Both perspectives are necessary, because spatial maps can only indicate where a given feature time series is expressed in the brain, but the cortical fingerprint of an event might be related to how the brain transitions through states, or critically, may differ across distinct kinds of events.

Here, we ask how perceived event boundaries relate to stimulus features, ongoing thought, and large-scale brain activity during movie-watching. We combine three independent samples who viewed the same eight movie clips, aligning these measures to a shared stimulus timeline: One marked perceived event boundaries, one reported ongoing thought using mDES, and one was scanned with fMRI, uninterrupted. We quantified visual, auditory, and semantic changes using pretrained neural networks (He et al., 2015; Hershey et al., 2017; Reimers & Gurevych, 2019), and we characterize neural activity both as whole-brain maps, and as moment-to-moment projections into gradient space (Margulies et al., 2016). We answered four primary research questions. First, we probe whether consensus event boundaries identify real transitions, by testing whether thoughts and features are more alike within-relative to between events. Second, we tested whether the aggregate brain states for features, thoughts, and events map onto a shared macroscale cortical architecture. Third, we tested whether boundaries reorient attention back toward the movie, by reducing distraction and a shift in cortical states toward sensory processing. Last, we tested whether boundaries were heterogeneous, with shared and distinct thought and cortical motifs across boundary types.

## Results

### Determining Moments of Consensus Event Boundaries

The first step in our analysis was to establish the consensus event boundaries, defined as local peaks in the smoothed boundary density exceeding the 90th percentile of that movie clip’s density (Michelmann et al., 2023). This yielded 64 event boundaries that fell within the temporal range covered by the available mDES probes (Figure 2A). The median interval between consecutive retained boundaries was 42.5 s (*M* = 47.8 s; *SD* = 34.2 s).

**Figure 2.**
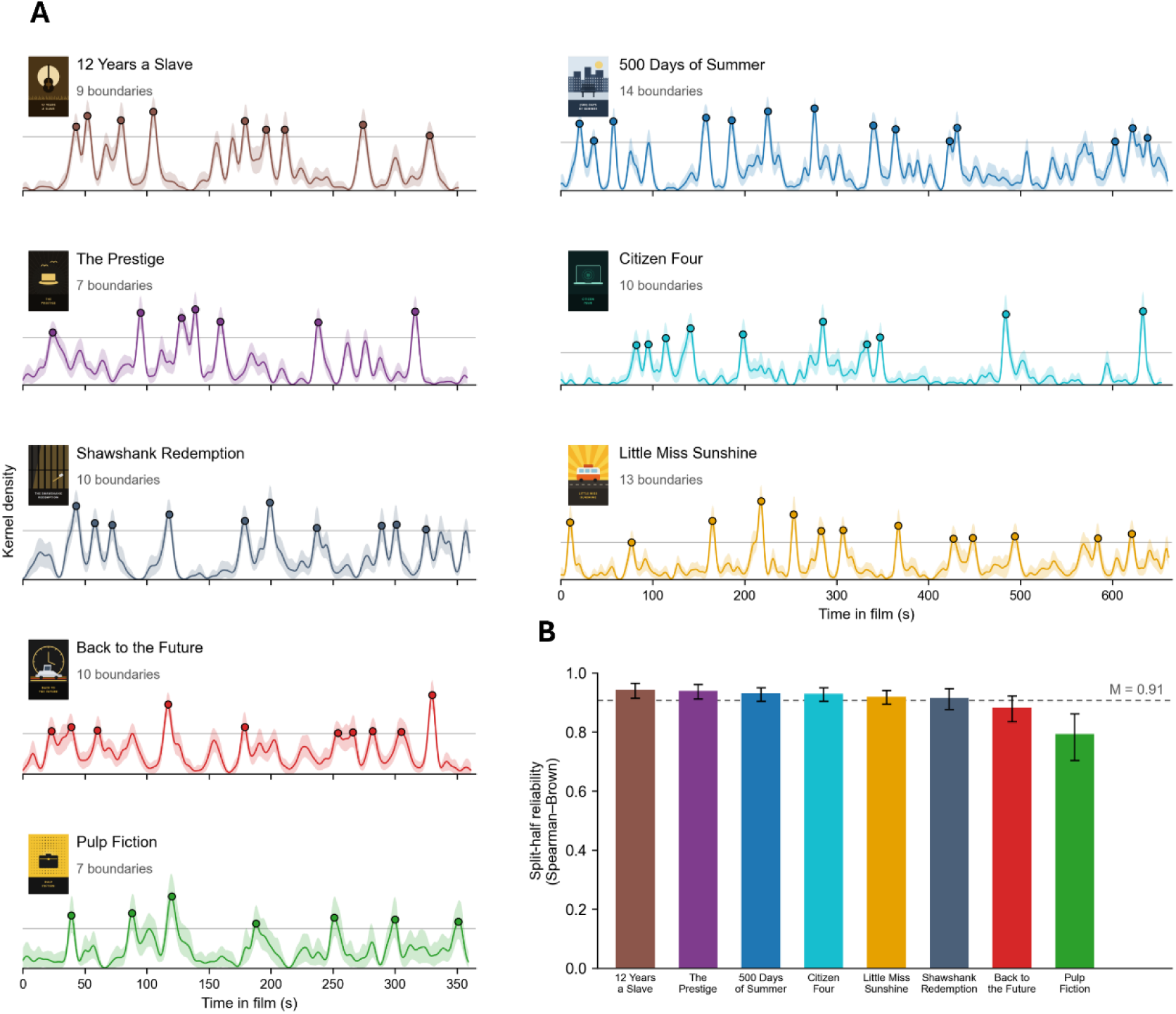
Per-movie consensus event boundaries and split-half reliability. Results of a kernel density estimation used to derive per-movie clip consensus boundaries. (A) Timestamps of button presses (n = 33 – 35, depending on movie) were smoothed with a Gaussian kernel (SD = 2.5 s) after removing repeat presses within 0.5 s. Peaks exceeding the 90th percentile (grey line) of the resulting density function were taken as consensus boundaries (coloured circles), 64 of which overlapped with periods of the movies where we had available multidimensional experience sampling (mDES) data from the thought sample. Shaded bands represent 95% confidence intervals from a participant-level bootstrap (2,000 resamples) of the kernel density. (B) Participants were split at random into halves, and a correlation was computed between the kernel density time course for each half then Spearman-Brown corrected. Bars are the mean over splits; error bars are the 2.5-97.5 percentiles of the resampling distribution, and the dashed line is the mean across films.

We next assessed how reliably participants identified the same moments as event boundaries using a split-half procedure. Across 2,000 random splits of participants within each movie, independently estimated boundary-density time courses were highly consistent. Spearman-Brown-corrected reliability ranged from .79 to .94 across movies (*M* = .91; Figure 2B). Pulp Fiction had the lowest reliability, likely because it also contained the fewest boundary responses (Supplementary Table 3).

### Generating the Thought-Space

Having established the reliability of the identified event boundaries, we next examined the thought patterns that emerged while participants viewed the movie clips. We collected 3,741 probe responses from 226 participants. Following prior work using mDES (Chitiz et al., 2026; Chiou et al., 2026; Hardikar et al., 2024; Konu et al., 2021; Mckeown et al., 2023; Mckeown et al., 2021; Medea et al., 2018; Mulholland et al., 2025; Mulholland et al., 2023; Lei et al., 2026), we standardized responses to the 16 experience-sampling dimensions and applied PCA with varimax rotation to identify dominant thought patterns across probes. This yielded five components (Figure 3A), which together accounted for 55.6% of the variance in mDES responses^1^.

**Figure 3.**
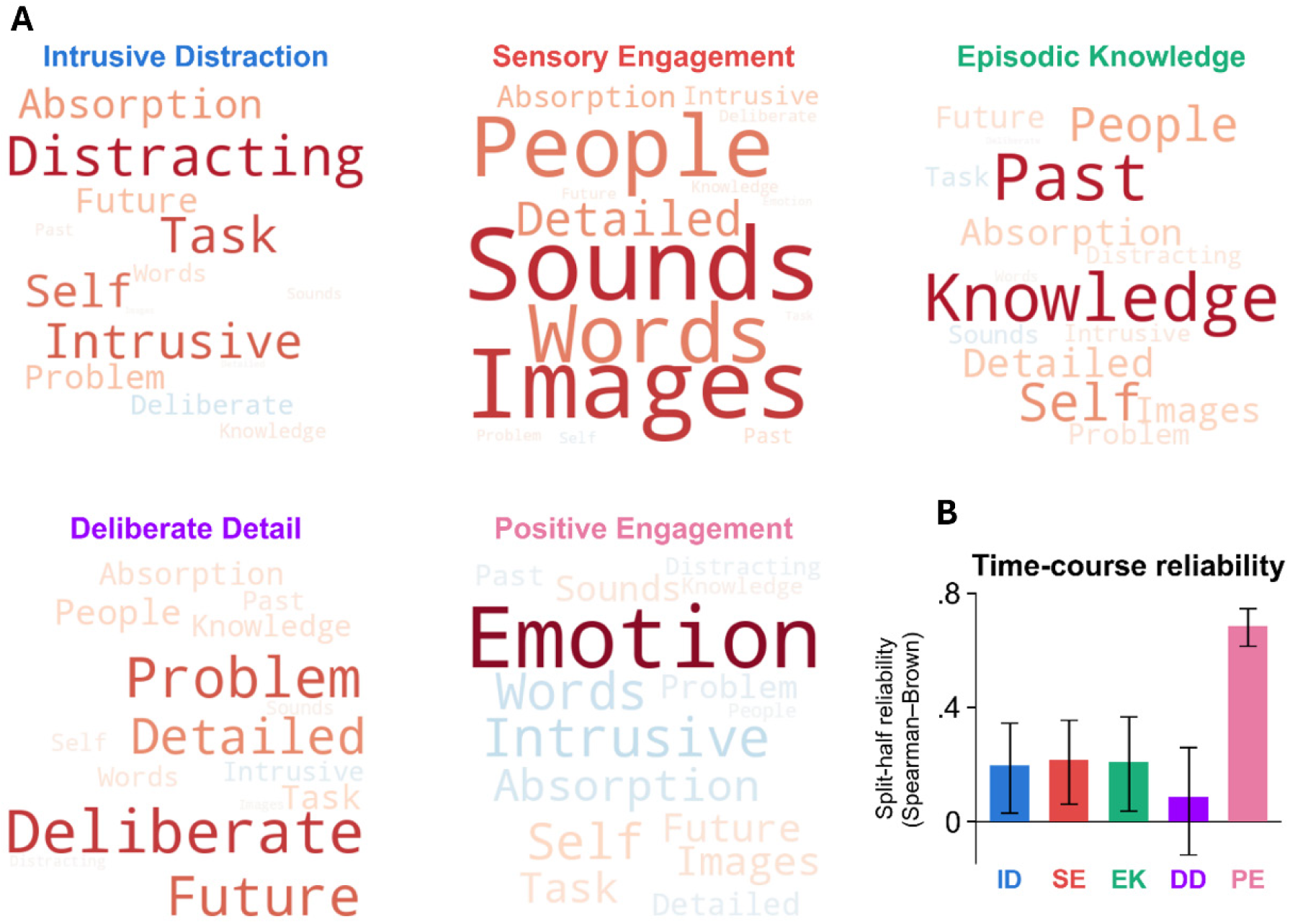
Word clouds representing the 5-component PCA solution and their reliability. Results of a principal components analysis which decomposed the 16 mDES items into low-dimensional components. (A) Word clouds representing the 5 retained components as thought patterns. Font size indicates the relative magnitude of the loading (bigger word = larger loading), and colour indicates the direction of the loading (red = positive, blue = negative). (B) Split-half reliability of the PC1-PC5 time courses. We randomly split participants in half, then correlated the halves and applied a Spearman-Brown correction. We repeated this process over 2,000 splits. Bars are the mean over splits; error bars are the 2.5-97.5 percentiles of the resampling distribution. Four of the five component time courses showed a consistently positive correlation across halves.

For ease of interpretation, we labelled each thought pattern according to the dimensions that loaded most strongly onto it. The first component, *Intrusive Distraction* (13.6% of variance), was characterized primarily by high loadings on the Distracting, Task, Intrusive, Self, and Absorption dimensions. The second, *Sensory Engagement* (12.9%), was characterized by Sounds, Images, People, Words, and Detailed. Using the same approach, we labelled the remaining components *Episodic Knowledge* (12.0%), *Deliberate Detail* (10.2%), and *Positive Engagement* (6.9%). Figure 3 depicts the loading structure of each thought pattern, with word size indicating loading magnitude and colour indicating loading direction. See Supplementary Table 4 for the rotated component matrix.

We next assessed whether this five-component structure could be reproduced in independent subsets of participants using bootstrapped split-half analyses. Across 2,000 resamples, the component solution showed high reproducibility (R-homologue = .96, 95% CI [.92, .99]; Tucker’s congruence = .93, [.87, .97]; subspace similarity = .91, [.83, .96]).

While this establishes that the thought patterns that emerge are reliable, we also aimed to determine how consistently they fluctuated in time for each movie. We conducted a second bootstrapped split-half reliability analysis examining whether the moment-to-moment component time courses, rather than the aggregate component structure itself, could be reproduced in independent sets of participants. Though these results were necessarily less stable than aggregated scores, split-half analyses indicated that four of the five component time courses were reliable: intrusive distraction (.20, 95% CI [.03, .34]), sensory engagement (.22, [.06, .35]), episodic knowledge (.21, [.04, .37]) and positive engagement (.68, [.61, .74]), with only the interval for deliberate detail falling below zero (.09, [−.12, .26]).

### Coherence of Event Structure

To answer our first research question, whether consensus event boundaries identify real transitions, we compared the similarity of moments within an event with that of moments separated by a boundary, for visual, auditory, and semantic embedding spaces and the thought space. There was significantly greater within- than between-event similarity in all four measures (Figure 4A; Supplementary Table 5). Visual embeddings were more strongly correlated within events than across adjacent events (.08 [.05, .13]), as were auditory embeddings (.03 [.02, .05]) and semantic embeddings (.02 [.006, .042]). Moments within an event were also closer together in the thought-space than moments separated by a boundary (.10 [.08, .13]). The effect was positive in all eight films for the thought-space, visual and auditory measures, and in seven of eight for the semantic measure (12 Years a Slave was the exception). To illustrate an example of a boundary crossing in each measure, we projected observations from either side of one consensus boundary into two dimensions using metric multidimensional scaling (see Figure 4B).

**Figure 4.**
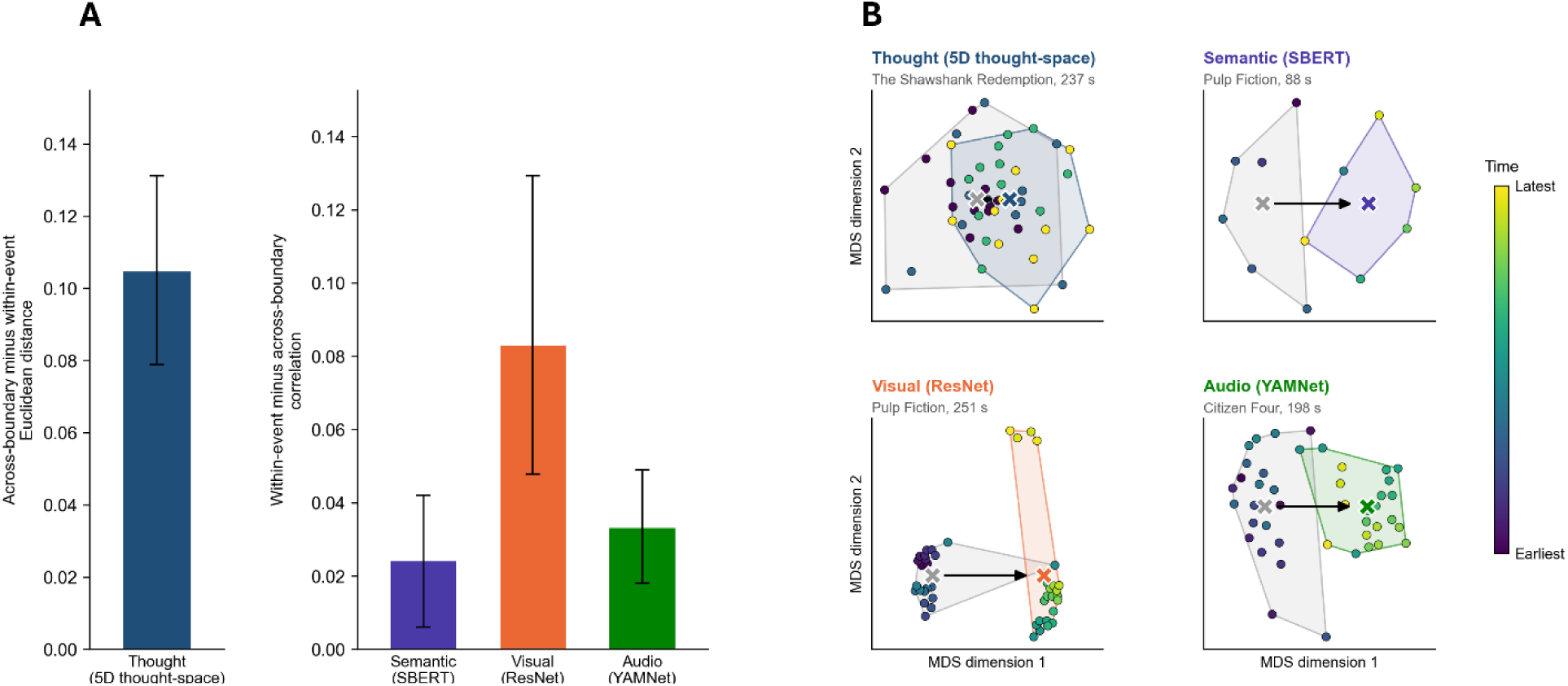
Consensus Boundaries Reliably Track Changes in the Thought-Space and Embedding Spaces. Results of a within-event versus across-boundary similarity analysis for each measure. (A) Bars show the difference between within-event and across-boundary similarity, with 95% confidence intervals from a movie-level cluster bootstrap of 2,000 resamples. We tested all four measures using the same 64 consensus boundaries. The test statistic for similarity in visual, auditory, and semantic features (right) is the mean correlation between embeddings, and the bar is within-event minus across-boundary correlation. For the thought-space (left) the measure is the Euclidean distance between moments in the five-dimensional component space, and the bar is across-boundary minus within-event distance. (B) An example of a single boundary crossing in each measure. For each measure, we used metric multidimensional scaling to project the embeddings or five-dimensional component scores for the time points within 21 seconds on either side of one consensus boundary into two dimensions. Component scores were taken at original probe times, with no interpolation. Each panel shows a different boundary, named above it, selected as a representative instance of that measure’s effect. Each point is one time point (or, for thought, one participant’s response at a probe), coloured by time within the window from earliest (dark purple) to latest (yellow). Shaded regions enclose the time points belonging to the event before the boundary (grey) and the event after it (coloured). Crosses mark the centroid of each event.

### Whole-Brain Analysis and Cortical Gradient Alignment

After establishing that event boundaries tracked changes in features and thought, we next tested our second research question, whether features, thoughts, and events map onto a shared macroscale cortical architecture. We fit whole-brain models with nine regressors (five thought patterns, consensus boundaries, and binarized visual, auditory and semantic change) at the participant-level, and combined them into a group-level mixed-effects analysis, yielding one group-level map for each regressor (Figure 5A; See Methods for more details), which can then be visualized in aggregate by computing the mean within the seven Yeo networks (Figure 5B; Supplementary Table 6).

**Figure 5.**
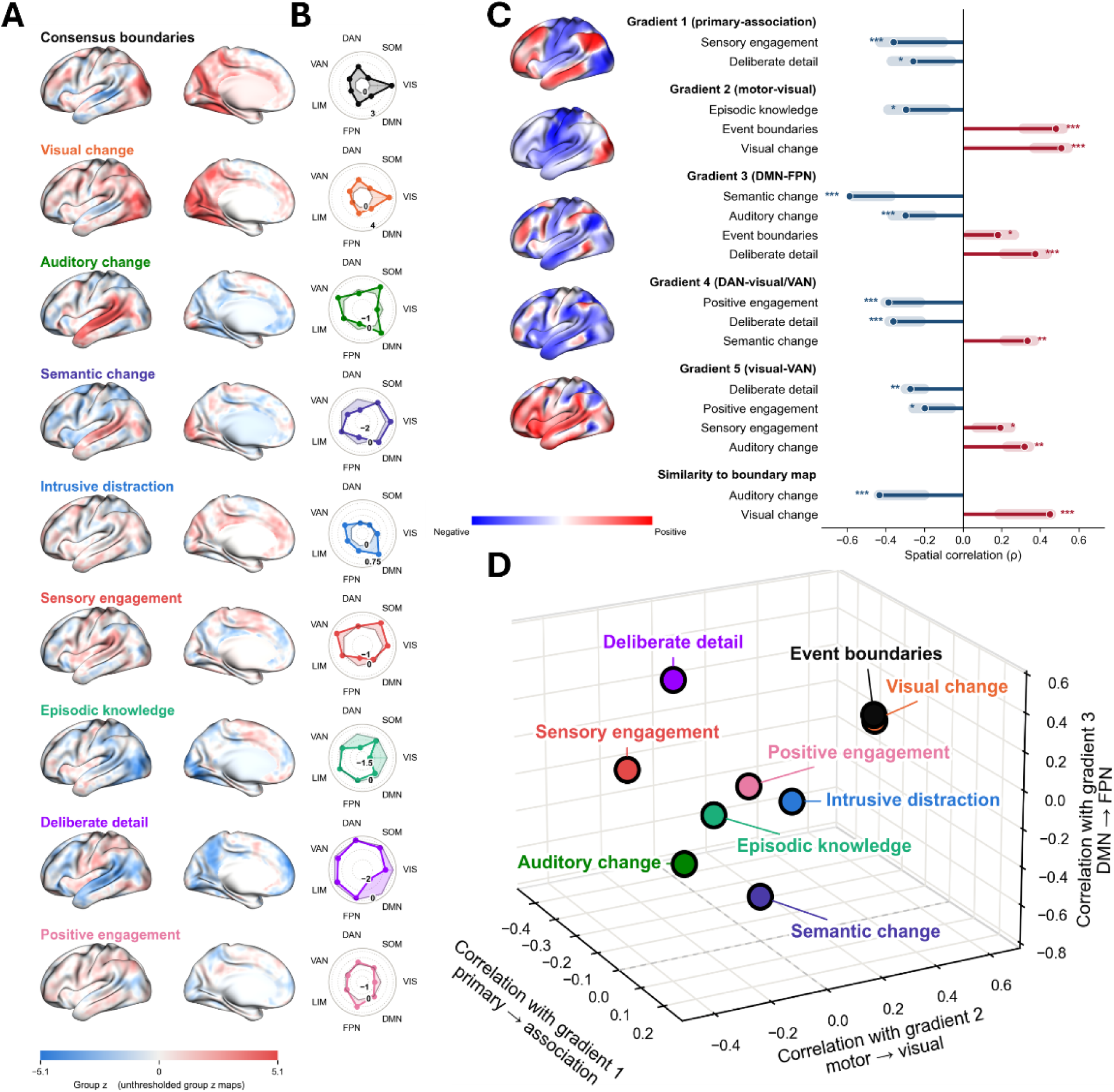
Whole-brain maps of boundaries, thoughts and features, and their positions in gradient space. Results of a general linear model predicting time courses of BOLD activity while participants viewed a selection of eight movies. (A) Left hemisphere lateral and medial views of whole-brain Z maps for each regressor entered into the model, including continuous interpolated thought pattern time courses and binarized consensus event boundaries, and features (visual, auditory and semantic). (B) Whole-brain maps visualized as aggregate scores according to a seven-network parcellation (Yeo et al., 2011), computed as the mean z statistic within each network. Values were first averaged within each parcel of the Schaefer 400-parcel atlas (Schaefer et al., 2018) and then across parcels belonging to each network, so that positive values indicate greater and negative values reduced activity. Each radar has its own radial scale. (C) Whole-brain maps were spin-tested against the five principal gradients derived from diffusion map embedding of resting-state data (Margulies et al., 2016) and against the whole-brain map for consensus event boundaries (bottom). Values are Spearman rank correlations between each unthresholded map and each gradient, or the consensus boundary map. Positive values indicate alignment with the positive end of a gradient (e.g. association cortex, for gradient 1) and negative values with the negative end (e.g. primary sensory cortex, for gradient 1). Comparisons are shown where the spin test was significant and the confidence interval excluded zero. Error bars represent the 95% confidence interval from 2,000 bootstrap resamples of the 74 runs. (D) Whole-brain maps from panel A projected into a three-dimensional scatter plot, based on their Spearman correlations with the first three gradients. The Gradients 1, 2, and 3 values provide X, Y, and Z coordinates for plotting, such that physical distance between points indicates similar correlations on the three gradients. Maps with *any* significant spin-test on *any* gradient are outlined in black.

Next, we situated each map within the first five cortical gradients (Margulies et al., 2016). Gradient 1 (g1) runs from primary sensory to association cortex; g2 from motor to visual cortex; g3 from the default mode network (DMN) to the frontoparietal network (FPN); g4 from the dorsal attention network (DAN) to visual cortex and the ventral attention network (VAN); and g5 from visual cortex to the VAN.

Sixteen of the map-by-gradient spatial comparisons exceeded their spin test nulls (Alexander-Bloch et al., 2018), with bootstrap resampled confidence intervals excluding zero, and two more were significant but these intervals did include zero (Sensory Engagement on g3; Intrusive Distraction on g4). The robust associations are depicted in Figure 5C, and all associations are reported in Supplementary Table 7A. Figure 5D shows each map’s position along the first three gradients, where similar states appear closer together in physical space (e.g. visual change and consensus boundaries).

Focusing on the thought patterns, Sensory Engagement sat toward the primary sensory end of g1 (*r_s_* = −.36, *p* < .001) and the VAN end of g5 (*r_s_* = .19, *p* = .021). Deliberate Detail sat toward the primary end of g1 (*r_s_* = −.26, *p* = .047), the FPN end of g3 (*r_s_* = .37, *p* < .001), the DAN end of g4 (*r_s_* = −.36, *p* < .001) and the visual end of g5 (*r_s_* = −.27, *p* = .003). Positive engagement loaded similarly on the latter two (g4 *r_s_* = −.39, *p* < .001; g5 *r_s_* = −.20, *p* = .025), and Episodic Knowledge was positioned away from visual processing, toward the motor end of g2 (*r_s_* = −.30, *p* = .015). Turning our attention to features and events, visual change and consensus boundaries both sat toward the visual end of g2 (*r_s_* = .51, *p* < .001 and *r_s_* = .48, *p* < .001, respectively), and event boundaries were also positioned toward the FPN end of g3 (*r_s_* = .18, *p* = .019). Semantic and auditory change were toward the DMN end of g3 (*r_s_* = -.59, *p* < .001; *r_s_* = -.30, *p* < .001). Semantic change in particular also pushed toward the visual/VAN end of g4 (*r_s_* = .33, *p* = .001), and auditory change toward the VAN end of g5 (*r_s_* = .32, *p* = .002). Generally, positioning these maps in gradient space revealed that external engagement and detail-oriented thoughts aligned with sensory, control, and goal-directed axes, while episodic thought was directed away from visual processing. Additionally, the feature maps were meaningfully separated by modality.

Finally, we examined how closely each thought and feature map correlated with the consensus boundaries map. It correlated positively with visual change (*r_s_* = .45, p < .001) and negatively with auditory change (*r_s_* = −.43, *p* < .001; Figure 5C), but not with semantic change (*r_s_* = −.13, *p* = .189) or with any thought pattern (all *p* ≥ .056; Supplementary Table 7B).

### Predicting Event Boundary Identifications

Our third research question, whether boundaries reorient attention back toward the movie, remained unclear based on the preceding analyses. While thoughts and features show reliable spatial associations with whole-brain cortical motifs, event boundary expression was only related to the maps corresponding to visual and auditory feature change. Most importantly, while events recruited both visual and FPN systems, they did not relate to any of the thought patterns, even deliberate detail, which shared their alignment with the FPN end of gradient 3. There are two possible explanations for this: One is that events are not a monolith, but instead are driven by different characteristics depending on the event, and may relate to different thought patterns, which are then obscured when aggregated. Alternatively, the relationship between events and thought may lie in the transition between brain states instead of the momentary position, which a time-collapsed map could not capture. These accounts may both be true, but both imply that the relationship between events and thought is invisible to static topography and requires time-resolved analysis.

Therefore, we next asked whether temporal dynamics in thoughts and features around event boundaries predicted whether participants identified those boundaries. Across the event segmentation sample, the gradient and feature models included 106,097 participant-seconds from 35 participants viewing eight films, 3,437 of which contained a boundary press; the thought-pattern model, fitted at probe resolution, included 8,365 participant-by-window observations, 2,927 with a press. We tested the three families of predictors separately using mixed-effects models and assessed significance against movie-wise circular-shift null distributions that preserved the temporal structure of the predictors.

At the level of thought patterns, events tended to be identified when there were lower levels of Intrusive Distraction (OR = 0.80, *z* = −2.37, *p* = .010), or (modestly) higher levels of Deliberate Detail (OR = 1.19, *z* = 1.82, *p* = .048). Positive Engagement, Sensory Engagement, and Episodic Knowledge did not predict event boundaries (all p ≥ .070; Figure 6A, top).

**Figure 6.**
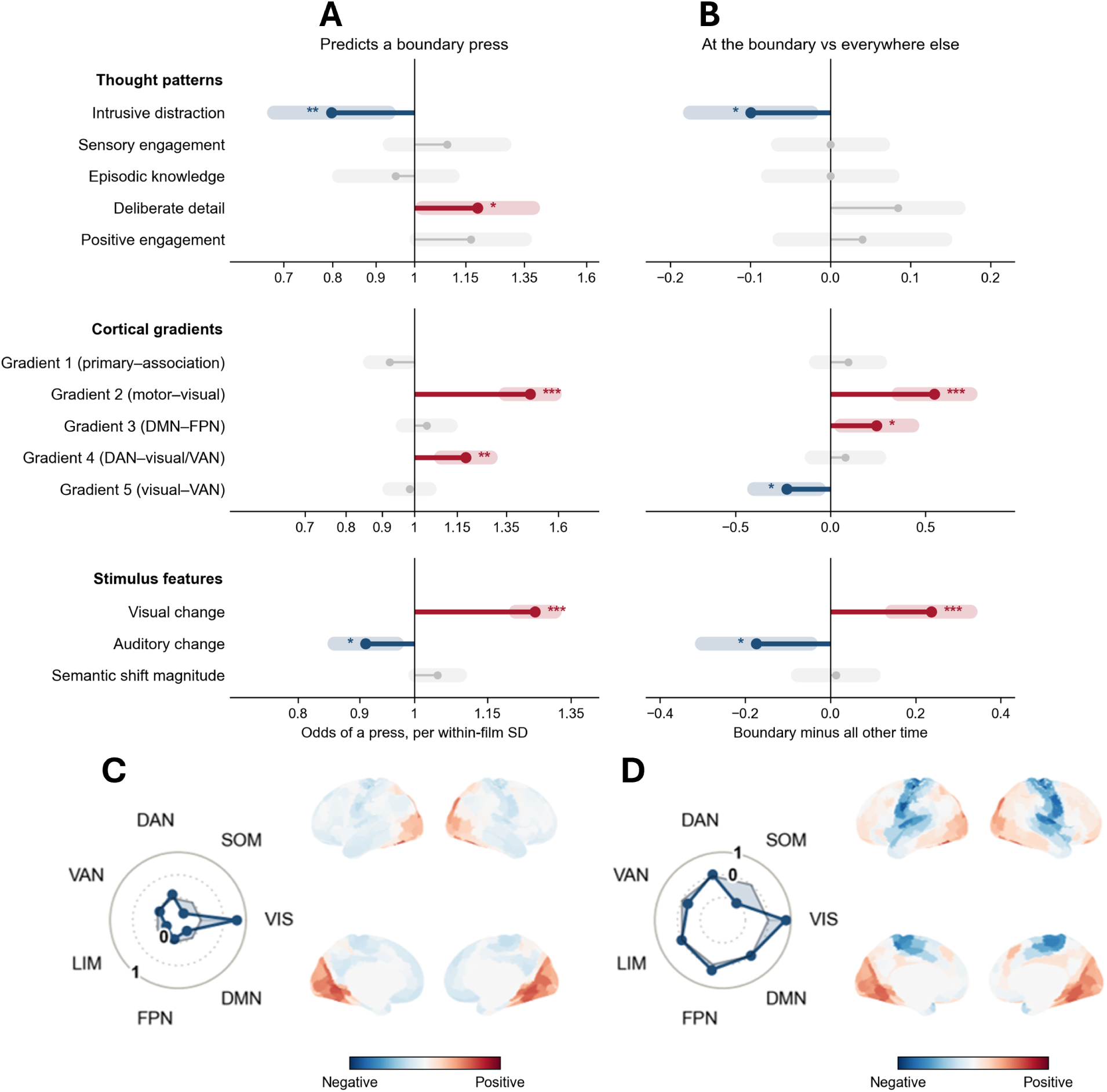
Gradient Position, Feature Change, and Thought Predict Boundary Presses; Gradient Position and Thought Change Across Boundaries. Results of three permutation-tested analyses relating the thought-space, the cortical gradients and feature change to boundary presses and to consensus boundaries. (A) What predicts a boundary press moment to moment. We modelled presses with binomial mixed-effects logistic regression with random intercepts for participant, film, and moment (the film-second for the gradients and features, the probe for the thought patterns), fitting the three predictor families in separate models and z-scoring all predictors within each film. Bars show the odds ratio per within-film standard deviation on a log scale. (B) The 7 s spanning each consensus boundary are compared with all other time points in the film. (C, D) For each model with a significant gradient effect, we averaged each of the five gradient maps within the 400 Schaefer parcels, weighted each parcellated map by that model’s coefficient for the corresponding gradient, and summed the five to produce one dot-product map per model. (C) The press model’s gradient coefficients, and (D) the across-boundary contrast’s gradient estimates. Surfaces show the model dot-product values on cortex, and radars show the mean of the same map over the parcels of each of the seven Yeo networks.

Boundary responses were more likely when brain states were positioned higher on gradients 2 (OR = 1.46, *z* = 11.25, *p* < .001) and 4 (OR = 1.18, *z* = 4.75, *p* = .002), indicating greater activation of visual cortex and VAN, as compared to motor cortex and DAN. The other gradients did not significantly relate to identified boundaries (all *p* ≥ .066; Figure 6A, middle). Boundary responses were also more likely during moments of greater visual change (OR = 1.26, *z* = 8.70, *p* < .001), and lower auditory change (OR = 0.91, *z* = -2.98, *p* = .015). Semantic shift magnitude did not significantly predict boundary responses (OR = 1.05, *z* = 1.44, *p* = .120; Figure 6A, bottom). See Supplementary Table 8 for the full table of results and coefficients.

### Boundary Crossing Analysis

To further clarify our third research question, whether boundaries reorient attention back toward the movie, we next probed for convergence by asking whether the same measures that predicted any identified event boundaries characterized the much more consistently identified consensus event boundaries. We examined whether measures differed at the moment of crossing a boundary relative to the rest of the movie.

Relative to all other moments in the movie (Figure 6B; Supplementary Table 9), crossing a boundary was associated with brain states positioned along the visual ends of both gradient 2 (*b* = 0.55, *p* < .001) and gradient 5 (*b* = −0.23, *p* = .022), and along the FPN end of gradient 3 (*b* = 0.24, *p* = .035). Not surprisingly, time periods spanning boundaries were also associated with greater visual change across boundaries (*b* = 0.24, *p* < .001), and less auditory change (*b* = −0.17, *p* = .018).

Among the thought patterns, only Intrusive Distraction showed a decrease across boundaries (*b* = −0.10, *p* = .030), while Deliberate Detail showed a marginal increase that did not reach significance (*b* = 0.08, *p* = .052). Gradients 1 and 4, semantics, and the remaining thought patterns did not differ (all *p* ≥ .39). For both this, and the previous analysis, predictive relationships hold using a leave-one-film-out analysis (see Supplementary Table 12); and are relatively insensitive to smaller shifts accounting for hemodynamic lag (lags of 4, 5, and 6 s are reported in Supplementary Table 13).

### Consensus boundaries fall into four subtypes

Strikingly, our univariate analyses indicated that the whole-brain states associated with aggregated event boundaries were not related to any of the thought patterns, despite aligning with the visual end of gradient 2 and the FPN end of gradient 3. There were also relatively modest changes related to aggregated events. This is likely because events can be related to changes of many different kinds, some of which are captured by our neural network feature time courses (see Figure 4).

Therefore, we aimed to not only analyze the nature of change in thought, feature and brain state across aggregated event boundaries, but also to identify specific subtypes of events with potentially different neural mechanisms. This was our fourth research question, whether boundaries were heterogeneous, with shared and distinct thought and cortical motifs across boundary types. To test this, we clustered the consensus boundaries by feature type. *k*-means clustering of the within-movie standardized dissimilarities selected four boundary types. We considered k from 2 to 6, since with 64 boundaries, larger *k* splits off clusters too small to estimate type-specific boundary responses. The four-cluster solution had the highest silhouette score (.35) and the most stability across subsamples (adjusted Rand index = .79) in that range. Extending the range to *k* = 10 produced a comparable silhouette score (.36), but the ten-cluster solution produced event types with only 4 to 9 boundaries (median = 6.5), which are too few to interpret (see Supplementary Table 10). Therefore, we retained the 4-cluster solution for subsequent analyses.

Using *k* = 4, the types (Figure 7A, B) were (1) visual boundaries (*n* = 20), where visual change was more extreme than at a typical boundary (visual *z* = 0.54, 95% CI [0.25, 0.84], auditory *z* = −0.69, 95% CI [−0.97, −0.40]; semantic *z* = -1.04, 95% CI [-1.18, −0.91]), (2) Semantic boundaries (*n* = 19), where semantic change was more extreme (visual *z* = −0.12, 95% CI [−0.56, 0.33]; auditory *z* = −0.51, 95% CI [−0.66, −0.38]; semantic *z* = 0.93, 95% CI [0.74, 1.11]). (3) Interruption boundaries (*n* = 18), defined by greater than usual audio and semantic change (visual *z* = 0.15, 95% CI [−0.21, 0.48]; auditory *z* = 1.24, 95% CI [1.05, 1.44]; semantic *z* = 0.52, 95% CI [0.17, 0.83]), and (4) Narrative boundaries (*n* = 7), where overall change was less extreme than usual (visual *z* = -1.63, 95% CI [-2.01, -1.25]; auditory *z* = 0.16, 95% CI [−0.61, 0.92]; semantic *z* = -0.90, 95% CI [-1.20, −0.53]).

**Figure 7.**
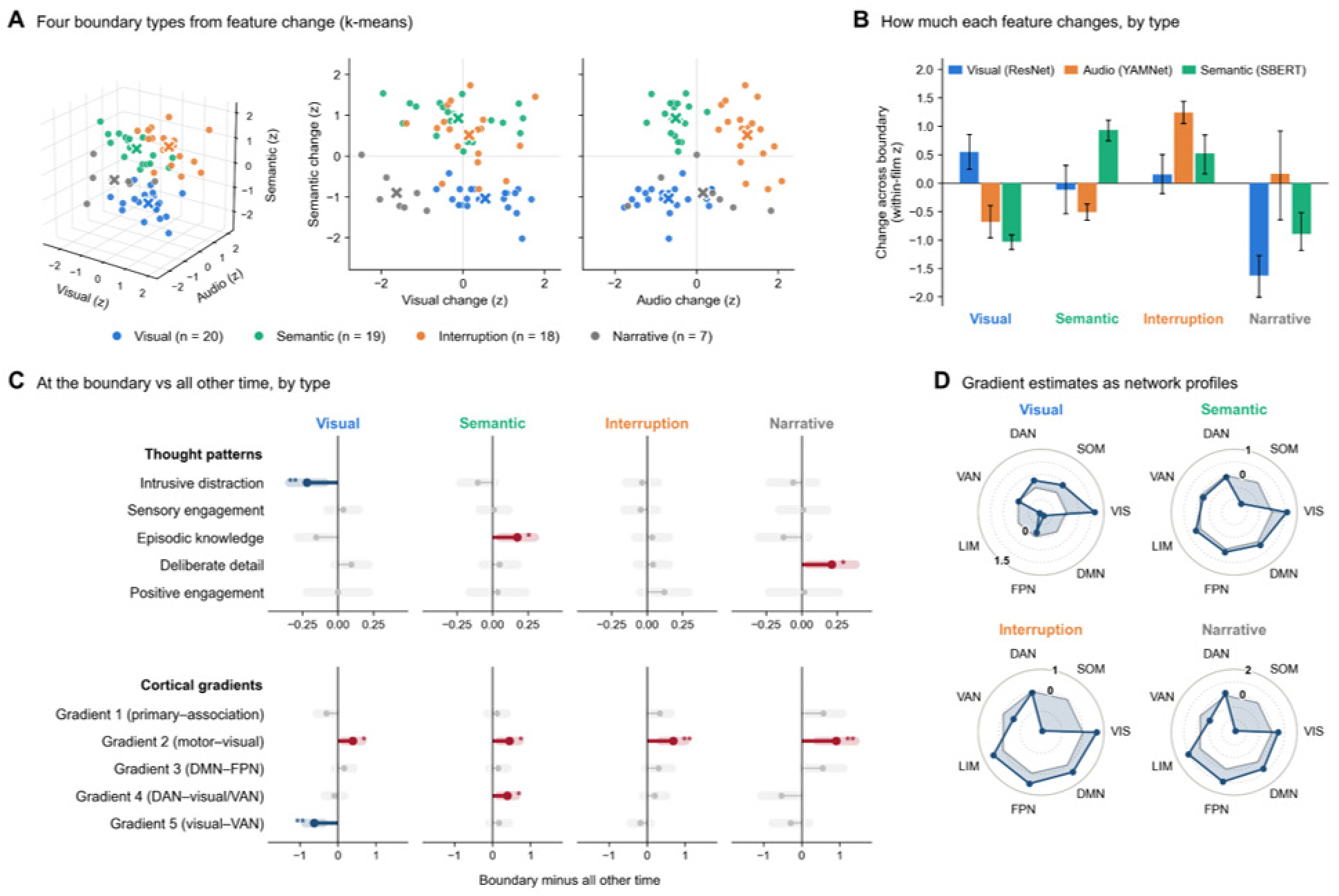
Boundary Types and the Across-Boundary Response Within Each Type. Results of a k-means clustering analysis to identify consensus boundary types based on feature changes. (A) Each of the 64 consensus boundaries plotted by its within-movie z-scored visual, auditory, and semantic change, in three dimensions (left) and in two planes sharing the semantic axis (right). Colours mark the four boundary types from k-means clustering and crosses mark each type’s centroid. (B) The four boundary types plotted based on how much the visual, auditory, and semantic content changed across each boundary, measured as the cross-boundary dissimilarity (1 minus similarity) of the feature vectors 5 s either side. We clustered these values, z-scored within each movie. Bars show the mean z-scored dissimilarity for each boundary type, and error bars are 95% bootstrap intervals over those boundaries (2,000 resamples). (C) The 7 s spanning the boundary compared with all other time in the movie. Shaded bands are 95% confidence intervals from circular-shift permutations (2,000 rotations, with boundaries rotated together with their type). (D) Radar plots depicting the gradient estimates of C expressed across seven Yeo networks for each type. To generate this, each gradient map was averaged within the 400 Schaefer parcels, weighted by that event type’s model estimates, and summed.

Critically, the four boundary types were comparable in their kernel density (0.006 to 0.007), so the subtypes were not driven by strength, prominence or consensus. Further, each movie had at least three of the four event subtypes, meaning that the cluster solution was not driven by different genres of film.

### Changes when crossing a boundary are distinct across event subtypes

There were clear commonalities in brain space states when boundaries were traversed across the subtypes. Not surprisingly, based on the aggregated analysis of all events together (see above; Figure 6), the temporal transition across every event boundary subtype was associated with a shift to the visual end of gradient 2 (visual *b* = 0.39, *p* = .036; semantic *b* = 0.45, *p* = .023; interruption *b* = 0.69, *p* = .001; narrative *b* = 0.91, *p* = .002). However, in only semantic boundaries, gradient 4 shifted toward its visual/VAN end (*b* = 0.39, *p* = .022), possibly reflecting an attentional reorienting in response to a salient change. In only visual boundaries, gradient 5 shifted toward its visual end (*b* = −0.63, *p* = .001). Together, these findings suggest that all boundaries involve a reconfiguration toward basic visual sensory processing, but changes driven by higher-order features like semantics also involve a shift toward VAN activation.

There were also critical differences in the role of thought patterns while traversing event boundaries. Intrusive Distraction was lower at visual boundaries relative to other moments in the same movie (*b* = −0.22, *p* = .009), suggesting that large visual changes may possibly break observers out of inattentive states. Semantic boundaries, where there were large changes in meaning, were exclusively associated with increases in thoughts related to Episodic Knowledge, which could possibly be associated with reviewing or reinstatement of prior experience when a dramatic contextual shift occurs in a movie (*b* = 0.17, *p* = .024). Last, narrative boundaries, where there was relatively modest change in all feature types, were exclusively associated with increases in thoughts involving Deliberate Detail (*b* = 0.21, *p* = .039). See Supplementary Table 11 for the full boundary type results.

## Discussion

Across three independent samples watching the same eight movie clips, we combined segmentation, experience-sampling, and fMRI data to examine transitions around event boundaries, as a function of movie content, ongoing thought, and brain activity. We found that the events that were identified marked legitimate transitions, as thought patterns, and visual, auditory and semantic content were more similar within a given event than between two events. While aggregate whole-brain maps connected event boundaries primarily to the sensory features of the stimulus, time-resolved analyses indicated that traversing a boundary was linked with a reduction in internally-directed intrusive distracting thoughts, and a reconfiguration of cortical states toward visual processing and cognitive control. Grouping boundaries by the nature of how the content changed revealed that while orienting toward visual processing was common, changes in thought and brain state depended on the type of boundary, and the kind of change that defined it.

The most consistent finding was a shift toward visual cortex around event boundaries, particularly along gradient 2. This was consistently apparent across the aggregate whole-brain maps, but also when considering the temporal relationship, via individual boundary presses, consensus boundaries and all event subtypes. This pattern even extended to the event subtype that was characterized by relatively little feature change. Together, these findings indicate that a reorientation toward perceptual processing is a general principle of how event boundaries constrain the brain. This links directly to evidence linking visual discontinuities to event segmentation in movies (Cutting et al., 2012), and shorter events to fluctuations in sensory cortex (Baldassano et al., 2017). Accompanying the shift toward visual cortex though, was a parallel shift toward the FPN, emergent both from the aggregate whole-brain analysis and the consensus event boundaries, reflecting attention on the movie (Wallace et al., 2025) and the control involved in event updating, rather than the event model itself, which has more often been linked to the DMN (Baldassano et al., 2017). For individual event identifications, there was also a shift toward the visual/VAN end of gradient 4, suggesting that in addition to cognitively shifting focus to visual processing, perceiving a transition involves reorienting attention toward salient stimuli (Corbetta & Shulman, 2002; Das et al., 2026; Vossel et al., 2014).

The measured thought patterns allow for a richer contextualization of event structure, quantifying how people experience these event boundaries. As boundaries are traversed, intrusive distraction is lower relative to other moments in the movie, but the subtype analysis revealed that this effect was mainly carried by visual boundaries, consistent with ideas from event segmentation theory, that boundaries elicit a failure of event model predictions, which requires refocusing and updating (Zacks & Swallow, 2007). At individual boundary presses, reduced distraction was accompanied by greater deliberate detail, though this finding was less consistent across analyses. However, together, these findings are consistent with longstanding findings that task-unrelated thought comes with a decoupling from the external world (Smallwood et al., 2016; Smallwood et al., 2021), so there are trade-offs between distraction and detail-oriented thinking. Together, these findings imply that event boundaries are sometimes a response to structural changes in the external world, that then trigger the engagement of cognitive control to reset attention, driving the brain from internally focused or distracted states toward more detailed, stimulus-oriented states.

Event structure clearly constrains cortical and thinking states, but none of the whole-brain maps for any of the thought patterns resembled the event boundary map. Because boundaries are not uniform and can mark different kinds of change (Cutting, 2014; Kurby & Zacks, 2008; Zacks et al., 2010), treating all boundaries equally explains why these relationships were obscured. When we grouped consensus boundaries into four types based on their visual, auditory, and semantic change (labelled *visual, interruption, semantic,* and *narrative*), all were associated with increased visual activity, but distinct from one another in the thinking states they elicited. Visual boundaries were the only subtype associated with lower intrusive distraction and a shift toward the visual end of gradient 5, consistent with accounts linking externally focused thought to greater coupling with sensory input (Smallwood et al., 2008; Wallace et al., 2025). Semantic boundaries were instead associated with greater episodic knowledge and a shift toward the visual/VAN end of gradient 4. Changes in the meaning of dialogue provide exactly the situation where the preceding context must be integrated to allow interpretation, and consistent with this shift to episodic knowledge, boundaries are known to elicit rapid reinstatement of prior brain activity patterns and greater hippocampal-cortical interaction (Barnett et al., 2024; Silva et al., 2019). The shift in gradient 4 is consistent with attentional reorienting, given that VAN regions respond to narrative suspense and reduced DMN activity in movies (Bezdek et al., 2017) and to boundaries in spoken stories (Herrmann et al., 2026). Together, these findings reflect viewers drawing on prior episodic knowledge to interpret the shift (Hahamy et al., 2023; Varga et al., 2025; Zacks & Swallow, 2007), consistent with the association between episodic knowledge and better movie comprehension (Wallace et al., 2025).

The remaining boundary types were more subtle. Interruption boundaries had no clear thought relationships. Narrative boundaries, which involved little measured feature change, were associated with greater deliberate detail, which may reflect shifts in a shared internal model, rather than shifts in the external stimulus that our embedding measures capture (Zacks & Swallow, 2007). The aggregate maps showing null results with thought patterns then, does not mean that thoughts flow unconstrained through event transitions. Instead, the nuanced features of each event type are attenuated when they are pooled, and considering the types is necessary to reveal changes in thought, and to a certain extent cortical, states.

### Open Questions

We measured event boundaries, thought, and brain activity in separate samples, sharing a mutual stimulus which allowed temporal alignment. While this enabled rich comparisons of relatively uninterrupted thought and brain activity, it precludes our ability to link event, thought and brain *within* individual viewers. Future work could more directly test these within-participant effects by collecting a subset of these measures from the same sample. Within our mDES sample, participants only viewed isolated clips from each film, meaning that the fMRI participants, who watched full films, had additional context for the scenes. It will be important to measure evolving thought patterns across longer-duration stimuli to gain a full understanding of how event structure constrains thinking states. In this mDES sample also, we probed thought only every 10 to 15 s across participants, meaning that the data were somewhat sparse. This left deliberate detail as the least reliable thought pattern, meaning that while theory-consistent, the results should be interpreted with caution. This can be solved in future studies by more densely sampling thoughts around previously identified event boundaries. Last, some boundary types had relatively few instances, leaving only seven narrative boundaries and therefore limiting the precision of their estimates. Long-form stimuli have the potential to solve this problem as well.

## Conclusion

Our findings link event boundaries identified by one group to changes in thought and large-scale brain state in independent groups watching the same films. Events have primarily been studied as stimulus properties within narratives or as state changes in the brain. Our findings suggest that event structure also shapes the nature of ongoing experience and thought, and that different event types elicit different forms of thought, reorienting from intrusive distraction for overt perceptual changes, and leaning on prior knowledge when meaning shifts, with corresponding distinct brain state changes. Event structure may therefore be better understood not as a unified response to change at different time scales, but as a set of state shifts that match the demands of the kind of change that defines each boundary.

## Methods

For this study, a total of 335 participants were included to examine how perceived event boundaries relate to stimulus features, ongoing thought, and large-scale brain activity during movie-watching. We used three independent samples, who viewed the same set of movie clips with different task demands and contexts: one reported ongoing thought using mDES during viewing (n = 226), one marked perceived event boundaries (n = 35), and one viewed the movies while being scanned using fMRI (n = 74). The latter sample was drawn from the Naturalistic Neuroimaging Database (NNDb), an existing publicly available dataset (Aliko et al., 2020). Figure 1 summarizes the design and the measures derived from each sample.

**Figure 1.**
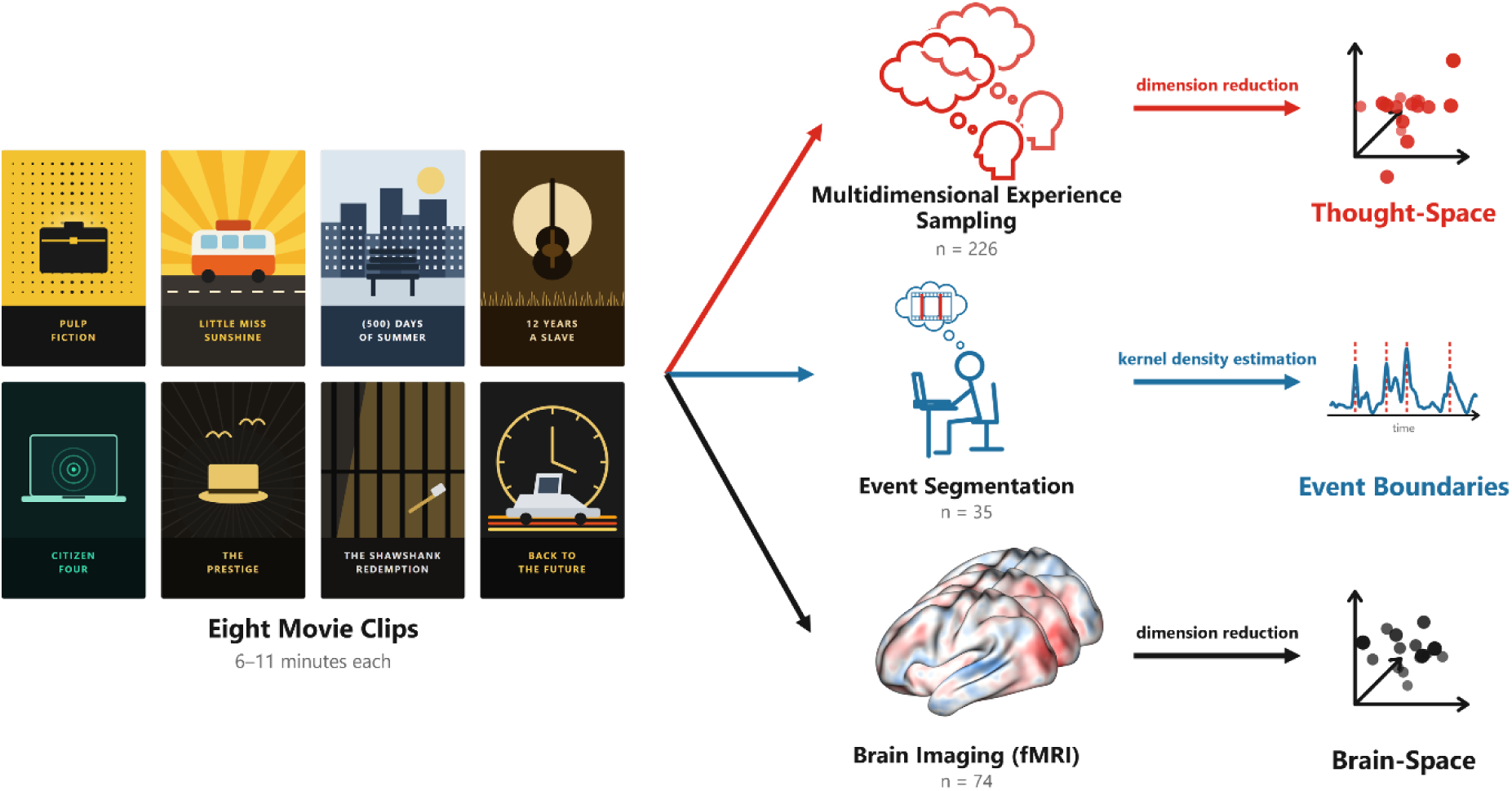
Study design. Three independent samples viewed the same eight movie clips (6–11 minutes each from Little Miss Sunshine, Pulp Fiction, 500 Days of Summer, 12 Years a Slave, Citizen Four, The Prestige, Shawshank Redemption, and Back to the Future), drawn from the Naturalistic Neuroimaging Database. The thought sample (n = 226) viewed a subset of either three or five movie clips and periodically scored their thoughts along 16 dimensions at counterbalanced time points throughout the clip. We decomposed these responses into a low-dimensional thought space using principal components analysis. The event sample (n = 35) viewed all eight clips and marked perceived event boundaries as they detected them. We applied kernel density estimation to derive consensus boundaries at the group level. The neuroimaging sample (n = 74) each viewed a single full-length film during fMRI, with coverage of all eight movies at the level of the entire sample. We extracted whole-brain activity corresponding to only the relevant clips and projected these data onto the first five cortical gradients (Margulies et al., 2016) to derive a low-dimensional brain space.

### Participants

#### Thought Sample

To measure experience using mDES, a total of 226 participants were recruited from the Queen’s University Department of Psychology participant pool between March 2023 and April 2024, in support of another project (see Wallace et al., 2026 for additional details). The sample included 184 women (81.4%), 34 men (15%), 6 non-binary (2.7%), 1 other gender identity (<1%), and 1 who preferred not to specify (<1%). The mean age was 19.03 years (*SD* = 1.30), ranging from 17 to 30 years.

#### Event Segmentation Sample

For event segmentation, we recruited 35 participants from the Queen’s University Department of Psychology participant pool, between October and November 2025. The sample included 30 women (85.71%), 4 men (11.43%), and 1 non-binary (2.86%). Their ages ranged from 17 to 34 years (*M* = 19.20, *SD* = 2.72). Most participants (34; 97%) resided in Canada and reported English as their primary language, while one participant from China reported Chinese as their primary language. The study received ethical approval from the Queen’s University General Research Ethics Board (#6044181). All participants provided informed consent and received a standard debriefing form upon completion.

#### Neuroimaging Sample

The neuroimaging sample was extracted from the Naturalistic Neuroimaging Database (NNDb; see Aliko et al., 2020 for additional details), which consisted of 74 participants (36 females, 38 males; mean age = 27.45 years, *SD* = 10.73; age range = 18-58 years, based on the 73 participants with available age data).

### Materials

#### Multidimensional Experience Sampling (mDES)

Participants in the thought sample reported their ongoing experience using mDES throughout the movie clips (see Supplementary Table 1) (Mulholland et al., 2023; Smallwood et al., 2021; Wallace et al., 2025; Wallace et al., 2026). mDES provides a detailed report of thought content by asking participants to rate their ongoing cognition along 16 dimensions (task, future, past, self, people, emotion, distracting, detailed, deliberate, problem, intrusive, knowledge, absorption, images, words, and sounds). For the emotion item, the scale ranges from 0 (“Negative”) to 10 (“Positive”). For the deliberate item, the scale ranges from 0 (“Spontaneous”) to 10 (“Deliberate”). For all other items, the scale ranges from 0 (“Not at all”) to 10 (“Completely”).

#### Stimuli

The movie clips consisted of 6-minute scenes from *Back to the Future, The Prestige, Pulp Fiction, 12 Years a Slave,* and *The Shawshank Redemption*; and 11-minute scenes from *Citizen Four, Little Miss Sunshine,* and *500 Days of Summer*. These films were selected based on their inclusion in the Naturalistic Neuroimaging Database (NNDb; Aliko et al., 2020), and their use in Wallace et al. (2026), where we derived the thought data from. The clips were originally chosen because they were relatively representative of the movie and were understandable as standalone scenes (i.e. without having seen the entire movie). For specific timestamps and brief scene descriptions, see Supplementary Table 2.

### Procedure

#### Multidimensional Experience Sampling

Participants attended a single laboratory session lasting approximately one hour. Participants viewed a random subset of 3 or 5 movie clips in randomized order. During each clip, viewing was interrupted at counterbalanced times to administer mDES probes. All participants provided informed consent and were compensated with either one course credit or $10.00 CAD. Across the entire sample, this results in 3,741 probes from 226 participants, with at least 11 participants responding at every probe time, covering every 10 seconds in the five shorter clips and every 15 seconds in the three longer clips. See the original manuscript (Wallace et al., 2026) for the full mDES procedure.

#### Event Segmentation

Participants attended a single laboratory session lasting approximately one hour. All participants first provided informed consent and completed a brief demographic questionnaire including their age, gender, sex, language, and country of residence. Participants completed the study individually on a laboratory computer while wearing headphones. Before viewing the movie clips, participants received brief written instructions to press the space bar whenever they perceived one meaningful event ending and another beginning (c.f. Jafarpour et al., 2022; Newtson & Engquist, 1976; Sargent et al., 2013; Sasmita & Swallow, 2023; Speer et al., 2003; Zacks & Swallow, 2007). These instructions were displayed on the laboratory computer to ensure consistency across participants. Participants first watched a 1-minute practice clip depicting an individual waking up and going through their morning routine (e.g., brushing their teeth, and putting on shoes), with clear event boundaries to familiarize themselves with the task. Participants then viewed each of the 8 experimental clips in random order and indicated perceived event boundaries in real time by pressing the space bar. After viewing all eight movie clips, participants received a standard debriefing form. Two participants left the study early, so 33 participants contributed data for Pulp Fiction and 34 for Little Miss Sunshine; all 35 contributed data for the other six clips.

#### Neuroimaging

fMRI data were acquired from the NNDb, an open-access database of preprocessed fMRI data in 3 mm MNI space (TR = 1 s) from participants who watched one of ten full-length films (Aliko et al., 2020). Twenty participants were recorded watching *500 Days of Summer*, eighteen watched *Citizen Four*, and six each of the remaining six films. We cropped each run to include only the TRs within the mDES probes’ coverage (261 TRs for the 6-minute clips and 586 TRs for the 11-minute clips). See the original manuscript for details about the acquisition and preprocessing procedures (Aliko et al., 2020).

### Neural Network Model and Layer Selection

We derived three feature change time courses, semantic, visual and auditory, using pretrained neural network models. In each case, we quantified change using cosine distance (1 minus cosine similarity) between temporally successive embeddings. For the visual and auditory change measure, this yields one value for each second of each film. For the semantic change measure, periods of silence are omitted, and therefore change is defined only at seconds when a new line of dialogue begins (see below).

#### Semantic

We extracted semantic features for transcribed dialogue using the Sentence-BERT all-MiniLM-L6-v2 model, trained to generate sentence-level embeddings (Reimers & Gurevych, 2019). Transcripts were generated using WhisperX in Python (Bain et al., 2023) and cross-referenced against the corresponding subtitle files (.srt) to identify and correct any transcription errors. We pulled embeddings, in the form of 384-dimensional vectors, for each transcript line, and calculated semantic change between successive lines as one minus cosine similarity (cosine distance; Zhuang et al., 2026) between the two corresponding vectors. These values were then assigned to the onset of the second line, so that a change spanning several seconds was counted only at the moment the change in dialogue occurs. If there were two or more lines in the same second, change from the preceding second was computed relative to the final line in that second, thereby capturing the total shift across lines. Seconds without line onsets were zero-coded and accounted for in subsequent analyses.

#### Visual

We extracted visual features for each frame using ResNet-50 (He et al., 2015) pretrained on ImageNet (Deng et al., 2009; torchvision IMAGENET1K_V2 weights). We sampled one frame per second, resized and centre-cropped each to 224 × 224 pixels, and normalized using the ImageNet channel means and standard deviations. Activations were read from the first residual block of the final convolutional stage (layer4.0) and averaged over the two spatial dimensions, providing one 2048-dimensional vector per second. Because categorical information in convolutional networks increases across the hierarchy and peaks in the final classification layers (Zeman et al., 2020), we chose layer4.0 to retain high-level scene and object structure while avoiding the most classification-specific (and therefore semantically driven) representations. Consistent with this, change in layer4.0’s embeddings showed a smaller correlation with our semantic change measure than the later ResNet-50 layers (see Supplementary Figure 1A).

#### Auditory

We represented the audio track from each movie clip using YAMNet, a MobileNet-architecture model trained on AudioSet (Gemmeke et al., 2017; Hershey et al., 2017). YAMNet is trained to analyze audio in fixed 0.96 s windows. By default, these windows are spaced 0.48 s apart, meaning that adjacent windows share overlapping audio. We doubled this default to prevent this overlap. We then averaged the depthwise and pointwise activations of layer 12 over time and frequency, concatenated them into a 1024-dimensional vector, and assigned each window to the second nearest its midpoint, averaging windows that fell in the same second. Layer 12 precedes the model’s final embedding and classification layers; representations at this depth capture acoustic structure while remaining distinct from explicit semantic categories (Giordano et al., 2023). Correlations with our measure of semantic change were small at every depth (mean *r* ≤ .11), including at layer 12 (*r* = .09; Supplementary Figure 1B), and fell further after layer 12, likely owing to the increasingly categorical representations of the final layers.

### Data Analysis

#### Event Boundary Identification

Participants in our event segmentation sample marked each time they perceived one meaningful event ending and another beginning while viewing eight movie clips (c.f. Jafarpour et al., 2022; Newtson & Engquist, 1976; Sargent et al., 2013; Sasmita & Swallow, 2023; Speer et al., 2003; Zacks & Swallow, 2007). To clean the data, we omitted any recordings that occurred within 0.5 seconds of each other. We then aggregated participant responses at the group level to produce a continuous time course of perceived boundaries for each movie clip. Next, we used Gaussian kernel smoothing (SD = 2.5 s) to estimate density of responses, and defined consensus event boundaries as local peaks in this density function that exceeded the 90th percentile of the density distribution for the corresponding movie clip. We then retained only consensus boundaries that fell within the window covered by the mDES probes.

We conducted a split-half analysis to assess the reliability of the estimated boundary-density time courses, separately for each movie clip. To do this, we randomly divided participants into two groups, independently estimated boundary-density time courses for each half and computed Spearman-Brown corrected correlations across the resulting time courses. This was repeated 2000 times to estimate the sampling distribution.

#### Thought Pattern Identification and Reliability

Thought patterns were identified from participants’ responses to the 16 mDES items. First, each item was standardized using a z-score transformation. Following previous applications of mDES (Chitiz et al., 2026; Chiou et al., 2026; Hardikar et al., 2024; Konu et al., 2021; Mckeown et al., 2023; Mckeown et al., 2021; Medea et al., 2018; Mulholland et al., 2025; Mulholland et al., 2023; Lei et al., 2026; Wallace et al., 2025), Principal Components Analysis (PCA) with varimax rotation was then applied to the standardized responses, and components with eigenvalues > 1 were retained to identify a low-dimensional set of recurring thought patterns, which together form the thought-space. Individual probes could then be expressed as component scores, which represented the extent to which a given probe response reflected each thought pattern.

We assessed the reliability of the resulting component structure using a bootstrapped split-half procedure (Chitiz et al., 2025; Mulholland et al., 2023). Across 2,000 iterations, we divided participants into independent halves and repeated the PCA separately within each half. Similarity between the resulting component solutions was quantified using three measures: R-homologue, which matches components from one half to its closest homologue in the other half, and computes the average correlation between their component scores; Tucker’s congruence, which computes the cosine similarity between matching components’ loading vectors; and subspace similarity, which asks whether the two halves span the same overall space of components, regardless of how the individual components are rotated or ordered within it.

We additionally assessed the *temporal* reliability of the five thought patterns across the movie clips. Across 2,000 bootstrapped split-half samples, participants were randomly assigned to two groups, and average component time-series were computed separately for each half. Participant-level component scores were mean-centred prior to aggregation. Corresponding time courses from the two halves were then correlated to quantify the extent to which moment-to-moment fluctuations in each thought pattern could be reproduced across independent groups of participants.

#### Examining Effects Across Event Boundaries

To test whether consensus boundaries marked genuine transitions, we asked whether time points falling within an event resembled one another more than time points separated by a boundary. For the same set of 64 consensus event boundaries, we tested for greater within- than between-event similarity in the five-dimensional thought-space, as well as the visual, auditory and semantic embedding spaces.

For the thought-space, we averaged component scores across participants at each probe time, linearly interpolated to 1 value per second to match the fMRI TR (1 s), and quantified dissimilarity between two time points as the Euclidean distance between their positions in the five-dimensional component space. The effect was calculated as across-boundary minus within-event Euclidean distance. For the three embedding spaces, we quantified similarity between two time points as the correlation between their embeddings, and the effect was calculated as within- event minus across-boundary correlation. Within-event comparisons for all measures were quantified by taking the average distance or correlation over all pairs of time points inside the same event, and between-event comparisons by averaging all pairs of time points in adjacent events that were separated by the event boundary. We computed the effect separately for each film, and inference was based on a movie-level cluster bootstrap in which we resampled the eight films with replacement 2,000 times.

#### Brain Analyses

To identify the neural correlates of event boundaries, feature changes, and thought patterns, we conducted whole-brain analyses using FSL’s FEAT framework. We ran a separate first-level FEAT analysis for each of the 74 participants in the neuroimaging sample, estimated in each participant’s native functional space (3 mm iso) and registered to a standard MNI152 template (2 mm iso). Because the existing fMRI data had already undergone standard preprocessing (Aliko et al., 2020), FEAT was used only for model fitting. Each model contained regressors for nine explanatory variables: five thought-patterns, consensus event boundaries, and visual, auditory, and semantic change. Thought-pattern scores were averaged across participants at each probe time and linearly interpolated to the temporal resolution of the fMRI data. Event boundaries were represented as a binary regressor marking consensus boundary locations. Visual, auditory, and semantic change regressors were binarized to identify the top 10% of change values within each movie clip (analogous to how consensus event boundaries were derived). For semantic change, we calculated this threshold across sentence-level observations rather than across all seconds. We additionally included a binary dialogue-presence regressor as a covariate to account for the presence versus absence of speech. All regressors were convolved with a double-gamma hemodynamic response function.

We then combined all the first-level estimates using FSL FLAME 1 mixed-effects modelling with a single group-level explanatory variable representing the group mean. This produced one group-level statistical map for each of the nine explanatory variables.

To characterize the resulting maps at the level of canonical functional networks, we summarized each unthresholded group map across the seven Yeo networks using the Schaefer 400-parcel cortical parcellation. For each network, we calculated the mean z statistic, first averaging within each parcel and then across the parcels belonging to that network, so that positive values indicate greater and negative values reduced activity.

We then characterized each group map’s position along the first five cortical gradients described by Margulies et al. (2016). We projected each group z map onto the fsLR 32k surface and calculated the Spearman rank correlation between the map and each gradient across all vertices. Because each gradient assigns every vertex a coordinate between its two ends (for example, from primary cortex to association cortex for gradient 1), a positive correlation indicates that the z statistic increases toward that gradient’s upper end and a negative correlation that it increases toward the lower end. The statistical significance of these positions was determined using spherical spin tests: each map was rotated 2,000 times (Alexander-Bloch et al., 2018), preserving its spatial autocorrelation, and the observed correlation was compared against the resulting null distribution. To describe the precision of each correlation, we calculated 95% bootstrap confidence intervals by resampling the 74 runs 2,000 times, rebuilding the group map from each resample and recomputing the correlation. We used the same spin test and bootstrap procedure to test the spatial correlation between each map and the consensus boundary map. For each pair, voxels were restricted to those present and finite in both maps, and a Spearman correlation was calculated between the two unthresholded *z* maps.

#### Generating the Cortical Brain Space

The preceding measures estimate aggregate whole-brain maps for each regressor, but abstract across moment-to-moment changes. The time course is likely to be more relevant in understanding changes as event boundaries emerge. To obtain a low-dimensional time course of brain activity, we adopted a “brain space” approach (Mckeown et al., 2023; Turnbull et al., 2020; Wallace et al., 2026) defined by the same five gradients from Margulies et al. (2016) shown in Figure 5C. We projected each movie into the cortical brain space using the group-averaged BOLD time series for that clip. At every acquisition (TR = 1 s), the Schaefer 400 parcellated activity map for that moment was correlated (Spearman rank) with each of the five gradient maps (Mckeown et al., 2025; Mckeown et al., 2023; Wallace et al., 2025). This procedure returns five values per time point: coordinates that locate a momentary brain state within each of the five gradients.

#### Predicting Identified Event Boundaries

With time courses for the five thought patterns, the three feature types, and the gradient scores, we examined how well these predicted the incidence of event boundaries. To do this, we modelled individual boundary responses from the event segmentation sample using binomial mixed-effects logistic regression. For each participant, we coded whether they identified an event boundary for every second of each movie clip. Models included random intercepts for participant, movie, and within-movie time point.

We examined features, gradients, and thought patterns as three separate predictor families and standardized each continuous predictor within each movie. The features model included visual, auditory, and semantic change. Because we defined semantic change only at sentence transitions, we modelled semantic shift magnitude separately from transition occurrence, including a binary sentence-transition indicator as a covariate. The cortical-gradient model included the first five cortical gradient time courses. The time courses were derived from the BOLD signal and temporally shifted gradient values by 6 seconds to account for hemodynamic delay before standardization and modelling. For the thought pattern model, data were analyzed at probe resolution rather than using the interpolated thought-pattern time courses. Each probe was assigned a temporal window extending halfway to the adjacent probes on either side, such that every point in the movie was assigned to its nearest probe without overlapping windows. For each participant and probe, the binarized outcome measure indicated whether the participant had registered a boundary within the corresponding window. We entered the five thought-pattern component scores simultaneously as predictors.

Statistical inference was based on movie-wise circular-shift permutation tests. For each permutation, we circularly shifted predictor time courses within each movie, disrupting their temporal correlation with boundary responses while preserving their in-measure temporal autocorrelation. Predictors belonging to the same model were shifted together to preserve their intercorrelations. We repeated this procedure 2,000 times for each predictor family and calculated permutation-based *p*-values by comparing the observed statistic with its corresponding null distribution.

#### Boundary Crossing Analyses

While the previous analyses were designed to establish if brain and thinking states coincide with the identification of event boundaries, critical state changes also occur as event boundaries are traversed (Baldassano et al., 2017; Geerligs et al., 2022; Zacks & Swallow, 2007). To test for these changes, we next examined how thought patterns, gradients, and features changed around all consensus event boundaries and later divided by event subtype (see below). We compared the average measure during periods spanning an event boundary with all the equivalent periods of time from the remainder of the movie, to determine whether a given measure changed as a boundary was traversed.

To select a boundary-crossing window, we elected to mirror the precision of the approach used to initially identify the consensus boundaries, by smoothing the time course with a Gaussian kernel at SD 2.5 s. Here then, and consistent with prior work using short symmetric windows (Lee & Chen, 2024; Zheng et al., 2022), we defined the period crossing the consensus boundary as including the 3 s on either side of it. To account for hemodynamic lag, we shifted gradient time courses by 6 s before applying the windows.

We fit models predicting whether a span included an event boundary or not, using gradients and features at the movie-second level, and including random intercepts for movie. Thought pattern predictors were each individual participant’s responses at the original probe times, without interpolation or averaging across participants; a probe belonged to a window when its time fell inside it, and these models included random intercepts for participant, movie, and within-movie probe time. Inferences were made by comparing to a permuted noise distribution, generated using movie-wise circular shifts of the consensus boundary locations across 2,000 permutations. For each permutation, we shifted boundary positions and their associated windows together within each movie, preserving the number and spacing of boundaries while disrupting their temporal alignment with the measured time courses.

#### Defining Boundary Types

To determine whether boundaries cluster into specific types, characterized by different types of feature change, we characterized each of the 64 boundaries by how much the movie’s visual, auditory, and semantic content changed across it. For each boundary, we took the 5 s immediately before it and the 5 s immediately after it, excluding the boundary second itself, and represented every second in these windows by its visual, auditory, and semantic embedding vectors. Within each of the three feature spaces, we then computed the Pearson correlation between every possible pair of pre-boundary and post-boundary time points (25 pairs), and averaged these correlations to get a single cross-boundary similarity for that boundary. This gives one cross-boundary similarity score per boundary, per feature space. We used 5 s on each side of the boundary for this analysis, a wider window than the 3 s crossing window used for the boundary crossing contrasts because the aim here is to characterize the content on each side rather than the response at the moment of crossing: a transcript line begins within 5 s of a boundary for 67% of boundaries, compared with 56% within 3 s, so the narrower window would leave the semantic measure uninformative for many boundaries, while 5 s remains local, never reaching a neighbouring boundary (the closest pair is 8 s apart).

The visual and auditory vectors were per-second ResNet and YAMNet embeddings, as described above, taken from 5 s on either side of a boundary. The semantic vector was the sentence embedding of the transcript line being spoken at that second, so a boundary with no new line within 5 s on either side received a semantic similarity of exactly 1. We first converted each similarity score into a dissimilarity score (1-similarity) and z-scored them. These three dissimilarities within each movie were then entered into k-means clustering.

We fit k-means for two to six clusters and chose the number of clusters by the silhouette score (how well separated the clusters are) and by stability (how consistently the same boundaries grouped together when we repeated the clustering on random 80% subsamples of the boundaries).

#### Boundary Crossing Analyses for Event Subtypes

We repeated the boundary-crossing contrast (see above) within each boundary type for gradients and thought patterns. Time points were assigned to their nearest boundary of any type, as before, so windows were unchanged, and an observation belonged to the window of a given type when its nearest boundary was of that type. For each event type, we compared the average measure when traversing a boundary of that type, with all equivalent time spans that did not include an event of *any* type, ensuring that other boundary types did not contaminate the baseline. As before, inferences were made using movie-wise circular shifts across 2,000 permutations, with each boundary’s label carried with it so that every permutation preserved the number and spacing of boundaries of each type.

## Acknowledgements

This work was supported by Natural Sciences and Engineering Research Council of Canada (NSERC) Discovery Grants awarded to author JDW (RGPIN-2020-05958) and JS (RGPIN-2023-03496); a New Frontiers in Research Fund (NFRF-2021-00183) awarded to authors JS and JDW; an NSERC Canada Graduate Scholarship-Doctoral (CGS D-601452-2025) awarded to RSW; and an NSERC Canada Graduate Research Scholarship-Master’s (CGRS M) awarded to SK.

## Footnotes

1 Three of the clips also included a probe after the clip had ended; these contributed to the PCA solution but were not included in the time courses. Excluding them from the PCA produced an essentially identical solution (Tucker’s congruence ≥ .998); the five components explained 55.3% of variance, compared with 55.6% in the reported solution.

