## Supplementary Materials for "Thoughts and cortical responses are shaped by the type of change at event boundaries"

#### Supplementary Table 1 - Multidimensional Experience Sampling Questions

| Dimension | Question | Scale Low (0) | Scale High (10) |
|---|---|---|---|
| Task | My thoughts were focused on an external task or activity | Not at all | Completely |
| Future | My thoughts involved future events: | Not at all | Completely |
| Past | My thoughts involved past events: | Not at all | Completely |
| Self | My thoughts involved myself: | Not at all | Completely |
| People | My thoughts involved other people: | Not at all | Completely |
| Emotion | The emotion of my thoughts was: | Negative | Positive |
| Images | My thoughts were in the form of images | Not at all | Completely |
| Sounds | My thoughts were in the form of sounds | Not at all | Completely |
| Words | My thoughts were in the form of words | Not at all | Completely |
| Detailed | My thoughts were detailed and specific: | Not at all | Completely |
| Deliberate | My thoughts were: | Spontaneous | Deliberate |
| Problem | I was thinking about solutions to problems (or goals): | Not at all | Completely |
| Intrusive | My thoughts were intrusive: | Not at all | Completely |
| Knowledge | My thoughts contained information I already knew (e.g., knowledge or memories) | Not at all | Completely |
| Absorption | I was absorbed in the contents of my thoughts | Not at all | Completely |
| Distracting | My thoughts were distracting me from what I am doing | Not at all | Completely |

*Note.* List of 16-item mDES questions administered at each probe.

#### Supplementary Table 2 - Movie Clip Descriptions and Timestamps

| Movie | Time in Full Movie | Description |
|---|---|---|
| *12 Years a Slave* | 0:58:41 – 1:04:54 | On a pre-Civil War plantation, an owner forces enslaved workers to dance at night, and his jealous wife attacks a young, enslaved woman. One man is later sent alone on an errand and stopped by a group of white men. |
| *500 Days of Summer* | 1:00:06 – 1:11:36 | A young man reconnects with his ex-girlfriend at a coworker's wedding. At her party, a split screen contrasts what he expected with reality: she is engaged. |
| *Back to the Future* | 0:36:46 – 0:43:02 | A teenager stranded in the 1950s watches his future father get bullied in a diner, then saves him from a car and is hit himself. |
| *Citizenfour* | 0:00:37 – 0:11:38 | Encrypted emails from an anonymous intelligence insider are read aloud. They alternate with interviews and testimony about government mass surveillance. |
| *Little Miss Sunshine* | 0:25:17 – 0:36:47 | A family on a road trip argues over breakfast. When their van breaks down, they must push-start it and jump in while it moves. |
| *The Prestige* | 1:32:23 – 1:39:00 | A stage magician's wife confronts him over his secrets and later takes her own life. His rival recruits help for a final show built around a vanishing machine. |
| *Pulp Fiction* | 2:20:15 – 2:26:30 | During a coffee-shop robbery, a hitman turns the tables on the robbers and talks everyone out of a gunfight. |
| *The Shawshank Redemption* | 1:51:04 – 1:57:20 | A prison warden discovers an inmate has escaped through a tunnel hidden behind a poster. A narrator explains how it was dug. |

*Note.* Timestamps and descriptions for each of the eight movie clips.

**Supplementary Table 3 - Consensus Event Boundaries and Reliability**

| Movie | Consensus boundaries within mDES range (all) | Button presses | Split-half r [95% CI] |
|---|---|---|---|
| *12 Years a Slave* | 6 (9) | 432 | .94 [.91, .96] |
| *500 Days of Summer* | 11 (14) | 933 | .93 [.90, .95] |
| *Back to the Future* | 7 (10) | 463 | .88 [.83, .92] |
| *Citizenfour* | 10 (10) | 485 | .93 [.90, .95] |
| *Little Miss Sunshine* | 12 (13) | 689 | .92 [.89, .94] |
| *Pulp Fiction* | 5 (7) | 256 | .79 [.70, .86] |
| *The Prestige* | 6 (7) | 472 | .94 [.91, .96] |

| Movie | Consensus boundaries within mDES range (all) | Button presses | Split-half r [95% CI] |
|---|---|---|---|
| *The Shawshank Redemption* | 7 (10) | 420 | .92 [.88, .95] |

*Note.* Boundary counts show the number within the mDES time range, followed in parentheses by the total identified in each movie. Button presses are the total number of participant boundary responses after removing repeat presses within 0.5 s. Split-half r is the Spearman–Brown-corrected correlation between boundary-density time courses (Gaussian kernel, SD = 2.5 s) from random participant halves over 2,000 splits. Intervals span the 2.5th to 97.5th percentiles. All $p < .001$.

#### Supplementary Table 4 - Varimax Rotated Component Solution

| mDES Item | Component 1 *Intrusive Distraction* | Component 2 *Sensory Engagement* | Component 3 *Episodic Knowledge* | Component 4 *Deliberate Detail* | Component 5 *Positive Engagement* |
|---|---|---|---|---|---|
| *Task* | .654 | .052 | -.135 | .193 | .150 |
| *Self* | .549 | -.089 | .457 | .089 | .205 |
| *People* | -.025 | .520 | .383 | .210 | -.045 |
| *Knowledge* | .111 | .100 | .811 | .154 | .061 |
| *Absorption* | .439 | .357 | .235 | .161 | -.169 |
| *Words* | .159 | .509 | .044 | .130 | -.216 |
| *Past* | .074 | .170 | .783 | .118 | -.079 |
| *Deliberate* | -.190 | .100 | .027 | .705 | .015 |
| *Sounds* | .071 | .745 | -.118 | .060 | .115 |
| *Future* | .305 | .094 | .168 | .585 | .137 |
| *Problem* | .292 | .093 | .147 | .619 | -.097 |
| *Distracting* | .787 | .004 | .117 | .034 | -.066 |
| *Images* | .031 | .704 | .219 | .026 | .146 |
| *Emotion* | -.023 | .050 | .023 | .006 | .906 |
| *Intrusive* | .604 | .256 | .114 | -.115 | -.243 |
| *Detailed* | .032 | .461 | .272 | .474 | -.087 |

*Note.* PCA of mDES responses (n = 3,741), with varimax rotation and Kaiser normalization.

#### Supplementary Table 5 - Within-Event Versus Across-Boundary Similarity

| Measure | Units | Within event | Across boundary | Effect [95% CI] |
|---|---|---|---|---|
| *Thought-space* | Euclidean distance | 0.568 | 0.673 | .105 [.079, .131] |
| *Visual (ResNet-50)* | Correlation | 0.791 | 0.708 | .083 [.048, .129] |

| Measure | Units | Within event | Across boundary | Effect [95% CI] |
|---|---|---|---|---|
| *Auditory (YAMNet)* | Correlation | 0.758 | 0.725 | .033 [.018, .049] |
| *Semantic (Sentence-BERT)* | Correlation | 0.194 | 0.170 | .024 [.006, .042] |

*Note.* Within-event pairs: all pairs of seconds inside the same event. Across-boundary pairs: all pairs between adjacent events (64 boundaries). Intervals from a movie-level cluster bootstrap (2,000 resamples of the eight films).

### Supplementary Table 6 - Whole-Brain Group Maps by Yeo Network

| Map | Visual | Somato-motor | Dorsal attention | Ventral attention | Limbic | Frontoparietal | Default mode |
|---|---|---|---|---|---|---|---|
| *Consensus boundaries* | 2.80 | 0.23 | 1.50 | 0.89 | 0.65 | 1.49 | 0.81 |
| *Visual change* | 3.22 | 0.30 | 1.55 | 0.76 | 0.17 | 1.20 | 0.76 |
| *Auditory change* | −0.58 | 0.82 | −0.37 | 0.65 | 0.02 | −0.59 | 0.92 |
| *Semantic change* | 0.64 | 0.17 | −1.44 | −0.83 | 0.01 | −0.88 | 0.41 |
| *Intrusive distraction* | 0.11 | −0.04 | −0.06 | 0.30 | 0.17 | 0.21 | 0.56 |
| *Sensory engagement* | 0.45 | 0.87 | −0.04 | 0.78 | 0.16 | −0.79 | −0.28 |
| *Episodic knowledge* | −1.38 | −0.23 | −1.15 | −0.18 | 0.05 | −0.13 | −0.37 |
| *Deliberate detail* | −0.76 | −0.24 | −0.03 | −0.35 | −0.28 | −0.21 | −1.86 |
| *Positive engagement* | −0.51 | −0.01 | 0.16 | −0.18 | −0.07 | 0.54 | 0.01 |

*Note.* Values are mean z from the unthresholded group maps (FLAME 1, 74 runs, one model with all nine regressors), averaged first within each of the 400 Schaefer parcels and then across the parcels of each of the seven Yeo networks.

### Supplementary Table 7 - Whole-Brain Correlations

#### A. Spin-tested correlations with each gradient

| Map | Gradient | $r_S$ | 95% CI | *p* |
|---|---|---|---|---|
| *Event boundaries* | Gradient 1 (primary–association) | −0.07 | [−0.23, 0.09] | .675 |
| | **Gradient 2 (motor–visual)** | **0.48** | **[0.29, 0.54]** | **< .001** |
| | **Gradient 3 (DMN–FPN)** | **0.18** | **[0.001, 0.29]** | **.019** |
| | Gradient 4 (DAN–visual/VAN) | 0.06 | [−0.05, 0.17] | .670 |
| | Gradient 5 (visual–VAN) | −0.13 | [−0.21, −0.04] | .319 |
| *Visual change* | Gradient 1 (primary–association) | −0.10 | [−0.26, 0.05] | .542 |

| Map | Gradient | $r_S$ | 95% CI | $p$ |
|---|---|---|---|---|
| | **Gradient 2 (motor–visual)** | **0.51** | **[0.34, 0.57]** | **< .001** |
| | Gradient 3 (DMN–FPN) | 0.12 | [−0.06, 0.23] | .168 |
| | Gradient 4 (DAN–visual/VAN) | 0.01 | [−0.07, 0.15] | .946 |
| | Gradient 5 (visual–VAN) | −0.16 | [−0.27, −0.03] | .199 |
| *Auditory change* | Gradient 1 (primary–association) | 0.04 | [−0.14, 0.24] | .794 |
| | Gradient 2 (motor–visual) | −0.31 | [−0.41, −0.10] | .085 |
| | **Gradient 3 (DMN–FPN)** | **−0.30** | **[−0.39, −0.13]** | **< .001** |
| | Gradient 4 (DAN–visual/VAN) | 0.15 | [−0.01, 0.29] | .288 |
| | **Gradient 5 (visual–VAN)** | **0.32** | **[0.20, 0.37]** | **.002** |
| *Semantic change* | Gradient 1 (primary–association) | 0.00 | [−0.20, 0.20] | .975 |
| | Gradient 2 (motor–visual) | 0.00 | [−0.16, 0.17] | .980 |
| | **Gradient 3 (DMN–FPN)** | **−0.59** | **[−0.60, −0.35]** | **< .001** |
| | **Gradient 4 (DAN–visual/VAN)** | **0.33** | **[0.19, 0.40]** | **.001** |
| | Gradient 5 (visual–VAN) | −0.04 | [−0.10, 0.06] | .710 |
| *Intrusive distraction* | Gradient 1 (primary–association) | 0.09 | [−0.19, 0.31] | .254 |
| | Gradient 2 (motor–visual) | 0.04 | [−0.19, 0.24] | .616 |
| | Gradient 3 (DMN–FPN) | −0.03 | [−0.24, 0.20] | .574 |
| | Gradient 4 (DAN–visual/VAN) | 0.19 | [−0.08, 0.35] | .006 |
| | Gradient 5 (visual–VAN) | 0.03 | [−0.12, 0.16] | .596 |
| *Sensory engagement* | **Gradient 1 (primary–association)** | **−0.36** | **[−0.46, −0.08]** | **< .001** |
| | Gradient 2 (motor–visual) | −0.20 | [−0.35, 0.06] | .093 |
| | Gradient 3 (DMN–FPN) | −0.16 | [−0.34, 0.11] | .041 |
| | Gradient 4 (DAN–visual/VAN) | 0.18 | [−0.03, 0.30] | .078 |
| | **Gradient 5 (visual–VAN)** | **0.19** | **[0.04, 0.27]** | **.021** |

| Map | Gradient | $r_S$ | 95% CI | $p$ |
|---|---|---|---|---|
| *Episodic knowledge* | Gradient 1 (primary–association) | 0.15 | [−0.04, 0.31] | .188 |
| | **Gradient 2 (motor–visual)** | **−0.30** | **[−0.41, −0.06]** | **.015** |
| | Gradient 3 (DMN–FPN) | 0.05 | [−0.13, 0.21] | .527 |
| | Gradient 4 (DAN–visual/VAN) | −0.03 | [−0.17, 0.11] | .824 |
| | Gradient 5 (visual–VAN) | −0.09 | [−0.17, 0.05] | .446 |
| *Deliberate detail* | **Gradient 1 (primary–association)** | **−0.26** | **[−0.40, −0.03]** | **.047** |
| | Gradient 2 (motor–visual) | −0.10 | [−0.25, 0.11] | .387 |
| | **Gradient 3 (DMN–FPN)** | **0.37** | **[0.18, 0.46]** | **< .001** |
| | **Gradient 4 (DAN–visual/VAN)** | **−0.36** | **[−0.41, −0.19]** | **< .001** |
| | **Gradient 5 (visual–VAN)** | **−0.27** | **[−0.32, −0.17]** | **.003** |
| *Positive engagement* | Gradient 1 (primary–association) | 0.13 | [−0.05, 0.29] | .254 |
| | Gradient 2 (motor–visual) | −0.16 | [−0.28, 0.03] | .251 |
| | Gradient 3 (DMN–FPN) | 0.14 | [−0.05, 0.29] | .075 |
| | **Gradient 4 (DAN–visual/VAN)** | **−0.39** | **[−0.43, −0.19]** | **< .001** |
| | **Gradient 5 (visual–VAN)** | **−0.20** | **[−0.28, −0.03]** | **.025** |

**B. Spin-tested correlations with the consensus boundary map**

| Map | $r_S$ | 95% CI | $p$ |
|---|---|---|---|
| ***Visual change*** | **.45** | **[.16, .48]** | **< .001** |
| ***Auditory change*** | **−.43** | **[−.46, −.17]** | **< .001** |
| *Semantic change* | −.13 | [−.29, .07] | .189 |
| *Intrusive distraction* | .01 | [−.20, .20] | .816 |
| *Sensory engagement* | −.14 | [−.30, .07] | .095 |
| *Episodic knowledge* | −.02 | [−.22, .15] | .881 |

| Map | $r_S$ | 95% CI | *p* |
|---|---|---|---|
| *Deliberate detail* | −.05 | [−.20, .13] | .553 |
| *Positive engagement* | −.19 | [−.30, .04] | .056 |

*Note.* Spearman correlations of aggregate whole-brain maps for each measure with (A) each of the five gradients and (B) the consensus boundaries map. Unthresholded group z maps projected to the fsLR 32k surface. *p* from 2,000 spherical spin tests and intervals from 2,000 bootstrap resamples of the 74 runs. Correlations with a significant *p*-value and intervals not crossing 0 are shown in bold. CI's are rounded to two decimals except when additional precision is needed to show whether an interval includes zero.

#### Supplementary Table 8 - Predictors of Boundary Presses

| Family | Predictor | *b* (log-odds) | *SE* | *OR* | *z* | *p* |
|---|---|---|---|---|---|---|
| **Thought patterns** | *Intrusive distraction* | −0.226 | 0.095 | 0.80 | −2.37 | .010 |
| | *Sensory engagement* | 0.089 | 0.097 | 1.09 | 0.92 | .307 |
| | *Episodic knowledge* | −0.051 | 0.094 | 0.95 | −0.55 | .585 |
| | *Deliberate detail* | 0.172 | 0.095 | 1.19 | 1.82 | .048 |
| | *Positive engagement* | 0.154 | 0.093 | 1.17 | 1.65 | .070 |
| **Cortical gradients** | *G1 (primary → association)* | −0.081 | 0.033 | 0.92 | −2.47 | .066 |
| | *G2 (motor → visual)* | 0.377 | 0.034 | 1.46 | 11.25 | < .001 |
| | *G3 (DMN → FPN)* | 0.040 | 0.036 | 1.04 | 1.13 | .458 |
| | *G4 (DAN → visual/VAN)* | 0.167 | 0.035 | 1.18 | 4.75 | .002 |
| | *G5 (visual → VAN)* | −0.016 | 0.034 | 0.98 | −0.47 | .719 |
| **Feature change** | *Visual* | 0.231 | 0.027 | 1.26 | 8.70 | < .001 |
| | *Auditory* | −0.093 | 0.031 | 0.91 | −2.98 | .015 |
| | *Semantic shift magnitude* | 0.044 | 0.031 | 1.05 | 1.44 | .120 |

*Note.* Binomial mixed-effects logistic regressions, one per family, predictors z-scored within film, so OR is per within-film SD. Random intercepts for participant, movie, and the film-second for the gradients and features, the probe for the thought patterns. Gradients advanced 6 s for hemodynamic lag and thought patterns measured at probe resolution. *p* from 2,000 movie-wise circular shifts of the predictors.

#### Supplementary Table 9 - Crossing a Consensus Boundary Versus Other Movie Moments

| Family | Measure | *b* | *SE* | *z* | *p* |
|---|---|---|---|---|---|
| **Thought patterns** | *Intrusive distraction* | −0.099 | 0.042 | −2.35 | .030 |
| | *Sensory engagement* | 0.000 | 0.040 | 0.00 | .976 |
| | *Episodic knowledge* | 0.000 | 0.045 | 0.00 | .956 |
| | *Deliberate detail* | 0.084 | 0.043 | 1.96 | .052 |
| | *Positive engagement* | 0.040 | 0.066 | 0.61 | .460 |
| **Cortical gradients** | *G1 (primary → association)* | 0.093 | 0.052 | 1.81 | .392 |
| | *G2 (motor → visual)* | 0.547 | 0.051 | 10.81 | < .001 |

| Family | Measure | *b* | *SE* | *z* | *p* |
|---|---|---|---|---|---|
| | *G3 (DMN → FPN)* | 0.243 | 0.051 | 4.72 | .035 |
| | *G4 (DAN → visual/VAN)* | 0.079 | 0.052 | 1.53 | .474 |
| | *G5 (visual → VAN)* | −0.231 | 0.052 | −4.49 | .022 |
| **Feature change** | *Visual* | 0.237 | 0.050 | 4.76 | < .001 |
| | *Auditory* | −0.174 | 0.050 | −3.45 | .018 |
| | *Semantic shift magnitude* | 0.013 | 0.051 | 0.25 | .779 |

*Note.* The crossing window spans 3 s either side of each of the 64 consensus boundaries. Every other time point in the movie is the reference. Each time point belongs to its nearest boundary, so windows never overlap. b is the difference in z units. Gradients shifted 6 s for hemodynamic lag. Thought patterns measured via individual participants' responses at the original probe times, random intercepts for participant, movie and movie time. *p* from 2,000 movie-wise circular shifts of the boundaries.

### Supplementary Table 10 - K-Means Cluster Selection Criteria

| k | Silhouette | Stability (ARI) |
|---|---|---|
| *2* | .28 | .60 |
| *3* | .30 | .64 |
| *4* | .35 | .79 |
| *5* | .32 | .68 |
| *6* | .31 | .62 |
| *7* | .32 | .68 |
| *8* | .33 | .69 |
| *9* | .35 | .74 |
| *10* | .36 | .76 |

*Note.* Stability is the mean adjusted Rand index between the full solution and solutions on 500 random 80% subsamples of the 64 boundaries.

### Supplementary Table 11 - Within-Type Crossing Effects Relative to Nonboundary Moments

| *Family* | *Measure* | *Visual (20)* | *Semantic (19)* | *Interruption (18)* | *Narrative (7)* |
|---|---|---|---|---|---|
| **Thought patterns** | *Intrusive distraction* | **−0.22 (.009)** | −0.11 (.210) | −0.04 (.640) | −0.06 (.527) |
| | *Sensory engagement* | 0.04 (.616) | 0.01 (.921) | −0.05 (.582) | 0.01 (.873) |
| | *Episodic knowledge* | −0.15 (.058) | **0.17 (.024)** | 0.03 (.677) | −0.13 (.218) |
| | *Deliberate detail* | 0.09 (.214) | 0.05 (.490) | 0.04 (.622) | **0.21 (.039)** |
| | *Positive engagement* | 0.00 (.989) | 0.04 (.725) | 0.12 (.231) | 0.02 (.925) |
| **Cortical gradients** | *G1 (primary → association)* | −0.31 (.107) | 0.12 (.548) | 0.33 (.113) | 0.57 (.065) |
| | *G2 (motor → visual)* | **0.39 (.036)** | **0.45 (.023)** | **0.69 (.001)** | **0.91 (.002)** |

| *Family* | *Measure* | *Visual (20)* | *Semantic (19)* | *Interruption (18)* | *Narrative (7)* |
|---|---|---|---|---|---|
| | *G3 (DMN → FPN)* | 0.16 (.402) | 0.15 (.392) | 0.31 (.203) | 0.56 (.093) |
| | *G4 (DAN → visual/VAN)* | −0.09 (.605) | **0.39 (.022)** | 0.20 (.345) | −0.53 (.108) |
| | *G5 (visual → VAN)* | **−0.63 (.001)** | 0.17 (.378) | −0.17 (.373) | −0.29 (.337) |

*Note.* The difference between the 7 s crossing a boundary of that type and every time point not in any crossing window. Z scored. Parentheses in column headers indicate the number of boundaries per type. Units are $b(p)$. $p$ from 2,000 movie-wise circular shifts of the boundary positions with their type labels attached. Bold: $p < .05$.

### Supplementary Table 12 - Held-Out Movie Effects and Prediction Accuracy

| *Measure* | *Mean effect* | *Mean AUC* |
|---|---|---|
| **Predicting a boundary response** | | |
| *Intrusive distraction* | −0.23 | .546 |
| *Deliberate detail* | 0.17 | .540 |
| *Gradient 2 (motor–visual)* | 0.38 | .595 |
| *Gradient 4 (DAN–visual/VAN)* | 0.17 | .537 |
| *Visual change* | 0.23 | .557 |
| *Auditory change* | −0.09 | .530 |
| **Crossing vs rest: all boundaries** | | |
| *Intrusive distraction* | −0.10 | .532 |
| *Gradient 2 (motor–visual)* | 0.55 | .650 |
| *Gradient 3 (DMN–FPN)* | 0.24 | .583 |
| *Gradient 5 (visual–VAN)* | −0.23 | .559 |
| *Visual change* | 0.24 | .539 |
| *Auditory change* | −0.17 | .566 |
| **Crossing vs rest: visual boundaries** | | |
| *Intrusive distraction* | −0.22 | .571 |
| *Gradient 2 (motor–visual)* | 0.38 | .601 |
| *Gradient 5 (visual–VAN)* | −0.63 | .691 |
| **Crossing vs rest: semantic boundaries** | | |
| *Episodic knowledge* | 0.17 | .575 |
| *Gradient 2 (motor–visual)* | 0.45 | .589 |
| *Gradient 4 (DAN–visual/VAN)* | 0.39 | .591 |
| **Crossing vs rest: interruption boundaries** | | |
| *Gradient 2 (motor–visual)* | 0.69 | .652 |
| **Crossing vs rest: narrative boundaries** | | |
| *Deliberate detail* | 0.21 | .563 |

| *Measure* | *Mean effect* | *Mean AUC* |
|---|---|---|
| *Gradient 2 (motor–visual)* | 0.91 | .740 |

*Note.* Each analysis was refitted eight times, leaving out one film each time. Mean effect is the average of the eight training-set estimates (log-odds per within-film SD for presses, z-score difference for the crossing contrasts). Mean AUC is the mean, over held-out films, of how well the measure separates presses from non-presses, or crossing-window time points from the rest of the movie, in the film left out (.5 = chance).

### Supplementary Table 13 - Sensitivity of Gradient Results to Assumed Hemodynamic Lag

#### A. Predicting boundary presses: OR (p)

| Gradient | 4 s | 5 s | 6 s (reported) |
|---|---|---|---|
| *G1 (primary → association)* | 0.97 (.459) | 0.93 (.127) | 0.92 (.066) |
| *G2 (motor → visual)* | **1.24 (.001)** | **1.37 (< .001)** | **1.46 (< .001)** |
| *G3 (DMN → FPN)* | 1.10 (.060) | 1.07 (.187) | 1.04 (.458) |
| *G4 (DAN → visual/VAN)* | **1.14 (.018)** | **1.18 (.003)** | **1.18 (.002)** |
| *G5 (visual → VAN)* | 0.97 (.522) | 0.99 (.770) | 0.98 (.719) |

#### B. Crossing a boundary versus the rest of the movie: b (p)

| Boundaries | Gradient | 4 s | 5 s | 6 s (reported) |
|---|---|---|---|---|
| **All (64)** | *G1 (primary → association)* | 0.06 (.586) | 0.08 (.500) | 0.09 (.392) |
| | *G2 (motor → visual)* | **0.45 (< .001)** | **0.52 (< .001)** | **0.55 (< .001)** |
| | *G3 (DMN → FPN)* | **0.28 (.016)** | **0.27 (.023)** | **0.24 (.035)** |
| | *G4 (DAN → visual/VAN)* | 0.02 (.862) | 0.06 (.591) | 0.08 (.474) |
| | *G5 (visual → VAN)* | **−0.23 (.021)** | **−0.26 (.008)** | **−0.23 (.022)** |
| **Visual (20)** | *G1 (primary → association)* | −0.16 (.391) | −0.23 (.217) | −0.31 (.107) |
| | *G2 (motor → visual)* | 0.36 (.052) | **0.39 (.036)** | **0.39 (.036)** |
| | *G3 (DMN → FPN)* | 0.22 (.247) | 0.19 (.315) | 0.16 (.402) |
| | *G4 (DAN → visual/VAN)* | −0.09 (.627) | −0.08 (.645) | −0.09 (.605) |
| | *G5 (visual → VAN)* | **−0.57 (.002)** | **−0.62 (.001)** | **−0.63 (.001)** |
| **Semantic (19)** | *G1 (primary → association)* | 0.01 (.993) | 0.05 (.798) | 0.12 (.548) |
| | *G2 (motor → visual)* | **0.43 (.028)** | **0.47 (.016)** | **0.45 (.023)** |
| | *G3 (DMN → FPN)* | 0.23 (.187) | 0.20 (.248) | 0.15 (.392) |
| | *G4 (DAN → visual/VAN)* | 0.28 (.116) | **0.35 (.045)** | **0.39 (.022)** |
| | *G5 (visual → VAN)* | −0.02 (.920) | 0.05 (.811) | 0.17 (.378) |
| **Interruption (18)** | *G1 (primary → association)* | 0.18 (.386) | 0.25 (.236) | 0.33 (.113) |
| | *G2 (motor → visual)* | **0.52 (.023)** | **0.63 (.002)** | **0.69 (.001)** |
| | *G3 (DMN → FPN)* | 0.31 (.194) | 0.31 (.198) | 0.31 (.203) |
| | *G4 (DAN → visual/VAN)* | 0.09 (.664) | 0.15 (.485) | 0.20 (.345) |
| | *G5 (visual → VAN)* | −0.17 (.391) | −0.19 (.326) | −0.17 (.373) |

| Boundaries | Gradient | 4 s | 5 s | 6 s (reported) |
|---|---|---|---|---|
| **Narrative (7)** | *G1 (primary → association)* | 0.56 (.070) | 0.58 (.055) | 0.57 (.065) |
| | *G2 (motor → visual)* | **0.60 (.048)** | **0.78 (.012)** | **0.91 (.002)** |
| | *G3 (DMN → FPN)* | 0.53 (.103) | 0.55 (.091) | 0.56 (.093) |
| | *G4 (DAN → visual/VAN)* | −0.53 (.101) | −0.53 (.111) | −0.53 (.108) |
| | *G5 (visual → VAN)* | 0.00 (.998) | −0.17 (.573) | −0.29 (.337) |

*Note.* The press and all-boundary results held at lags of 4, 5 and 6 s. The gradient time courses were advanced by 4, 5 or 6 s before modelling. *p* from 2,000 movie-wise circular shifts. Bold: *p* < .05.

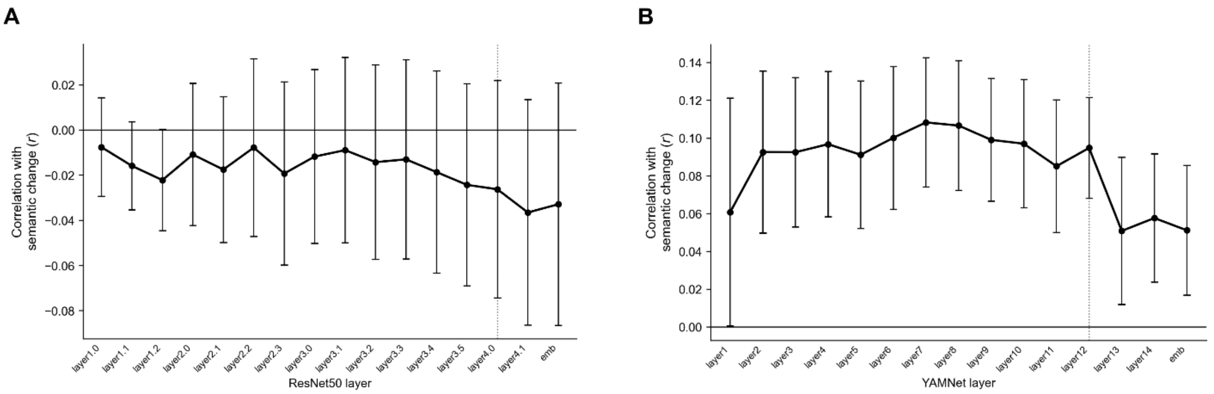


**Supplementary Figure 1. Correlations Between Model Layers and Semantic Change.** Each point is the correlation between change in a layer's features and semantic change, computed only over seconds containing dialogue onset, so that it reflects how much the meaning shifted rather than whether speech occurred. Points show the mean correlation across the eight films, and error bars show ±1 SE. (A) shows ResNet-50 (visual) and (B) shows YAMNet (auditory). Dotted lines mark the layers used as stimulus regressors (layer4.0 and layer 12), and "emb" is each network's final embedding.
